# Binocular integration of prey stimuli in the zebrafish visual system

**DOI:** 10.1101/2024.09.08.611846

**Authors:** Guangnan Tian, Thomas Ka Chung Lam, Gewei Yan, Yingzhu He, Biswadeep Khan, Jianan Y. Qu, Julie L. Semmelhack

**Affiliations:** Division of Life Science, The Hong Kong University of Science and Technology, Clear Water Bay, Kowloon, Hong Kong S.A.R., China; Department of Chemical and Biological Engineering, The Hong Kong University of Science and Technology, Clear Water Bay, Kowloon, Hong Kong S.A.R., China; Department of Electronic and Computer Engineering, The Hong Kong University of Science and Technology, Clear Water Bay, Kowloon, Hong Kong S.A.R., China

**Author notes:** Current address: Neuroengineering Laboratory, Brain Mind Institute & Interfaculty Institute of Bioengineering, EPFL, Lausanne, Switzerland. Current address: Department of Psychiatry and Behavioral Sciences, Stanford University, Stanford, CA, 94305, USA. These authors contributed equally: Guangnan Tian, Thomas Ka Chung Lam, Gewei Yan.

## Abstract

Most animals with two eyes combine the inputs to achieve binocular vision, which can serve numerous functions, and is particularly useful in hunting prey. However, the mechanisms by which visual information from the two eyes are combined remain largely unknown. Here, we designed a device to reversibly occlude the eyes of a head-fixed zebrafish larva, and used large-scale volumetric two-photon imaging to identify binocular neurons that respond to prey stimuli. We found these binocular prey-responsive neurons (bino-PRNs) were primarily located in three areas, the pretectum, thalamus, and nucleus isthmi. We then characterized the bino-PRNs’ functional properties, and found that their left and right eye receptive fields are offset to varying degrees, which would correspond to objects at naturalistic hunting distances for a larva with converged eyes. We also found that bino-PRNs have a significantly greater response in hunting trials, which could be the result of an eye convergence-related corollary discharge. We then optogenetically induced prey capture eye and tail movements, and found that this hunting command activates prey responsive neurons in the pretectum, thalamus, and nucleus isthmi. These findings indicate that bino-PRNs receive visual and motor input that would allow them to encode prey position in three dimensions.

## Introduction

All animals with binocular vision must integrate the two eyes’ signals to produce a unified percept and behavioral response, and this integration should occur in binocular neurons which receive input from both eyes. Binocular neurons in the visual cortex have been identified in non-human primates as well as mice, but it remains unclear where and how the inputs are first combined^1,2^. One proposed driver of the evolution of binocular vision is predation^3^, and binocular vision has been proposed to play a key role in hunting in a diverse range of animals, from insects to cephalopods to mice^4–6^. Mice have been shown to place prey objects in their binocular visual field, and ipsilateral retinal ganglion cell input is important for hunting^7^. However, it has been difficult to identify the binocular neurons that respond to the prey during hunting episodes, and to manipulate these neurons to determine their role in hunting behavior.

Binocular vision is also thought to play a role in zebrafish hunting behavior^8^. Zebrafish prey capture is typically initiated by the entry of a paramecium or other small moving object into the peripheral visual field. In response to the prey stimulus, the larva converges its eyes and hooks its tail in a J shape to turn toward the prey. These actions bring the prey from the peripheral to the central visual field,^9–13^ after which the larva may then perform a more symmetrical forward swim to approach the prey, before executing a capture strike to consume it^10,12^. Zebrafish larvae have laterally positioned eyes that normally have little binocular overlap, but the eye convergence at the beginning of a hunting episode creates a binocular zone^9^. Since eye convergence is a signature of hunting^9^, it is likely that binocular vision plays a key role in prey capture, although exactly how the larvae are using it remains unclear.

There are multiple possible functions for binocular vision: perhaps the simplest application is to increase the prey signal by summing two visual inputs, which could help larvae detect low-contrast prey^14^. A second possibility is that larvae use binocular vision for range-finding, or estimating the distance to prey, which could be achieved by foveating the prey with both eyes and triangulating the distance, if the extent of eye convergence is known^15^. A third option is that they are doing a more sophisticated form of stereopsis, and detecting binocular disparity, or the difference in the retinal position of the prey between the two eyes, to perceive depth^16^.

One way to clarify the role of binocular vision would be to find binocular neurons and study their functional properties. So far, although binocular interactions have been identified in the visual response to whole-field motion and luminance stimuli^17–19^, binocular prey-responsive neurons have not been found in zebrafish.

Here, we use large-scale imaging to identify binocular prey-responsive neurons (bino-PRNs) in multiple visual areas of the larval zebrafish brain. We find that these neurons are tuned to objects within striking distance of a hunting larva, and they also integrate eye convergence information, which could allow them to determine the location and distance of prey objects.

## Results

### Eye-blocker reversibly blocks input to one eye

To identify binocular prey-responsive neurons, we designed and 3D-printed a device to block one or both eyes selectively and reversibly by laterally moving its arms in front of the eyes (Figures 1A and 1B). The larvae were embedded in agarose with the tail free, and the eyes were kept in agarose to minimize large eye movements, which would change receptive fields of the neurons we image. Although the eyes were fixed, we could observe eye convergences of a few degrees in response to small, moving UV dots, which robustly trigger hunting behavior^20^. In order to restrict the prey object to a small region of the visual field, we designed a circulating stimulus which moved through an 8°×4° elliptical track, and presented this stimulus at a range of azimuths from −50° (left) to +50° (right) of the visual field (Figure 1B). We moved the eye-blocker to different positions between prey presentations such that all stimuli were presented during all four ocular conditions: left eye exposed (left-eyed), right eye exposed (right-eyed), both eyes exposed (binocular), and both eyes blocked (blocked) (Figure 1C). We tested the completeness of the block by recording visual responses in arborization field 7 (AF7), a retinorecipient region that responds to prey stimuli^21^, and found that, as expected, blocking one eye completely blocked the input to AF7 in the hemisphere contralateral to that eye (Figures 1D and 1E). This pattern was consistent across 7 fish (Figure 1F) and held true for all the stimulus azimuths (Figure 1G).

**Figure 1.**
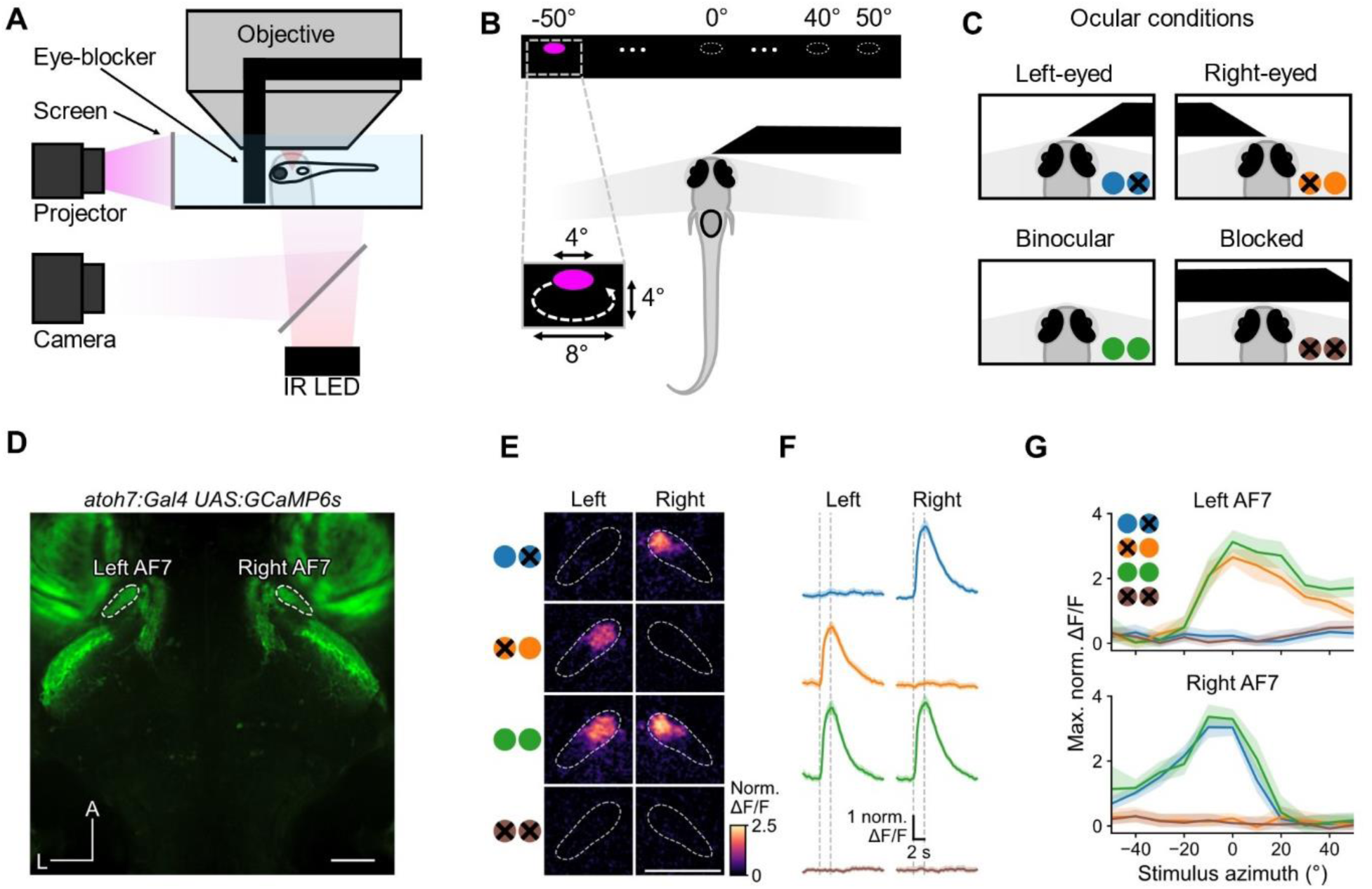
Reversible occlusion of one eye blocks the response to prey stimuli. (A) Schematic of the experimental setup: the eye-blocker is positioned in front of the fish to block visual input. (B) Bright UV circulating prey-like stimuli were presented at azimuths from −50° (left) to +50° (right). (C) Larvae were tested under 4 ocular conditions: right eye blocked, left eye blocked, neither eye blocked and both eyes blocked. (D) Baseline fluorescence image showing AF7s labelled with GCaMP6s (Scale bar, 50 μm). (E) Activation of AF7 in left eye, right eye, binocular, and fully blocked conditions in one example larva (Scale bar, 50 μm). Pixels are color-coded by trial-averaged increase in mean ΔF/F during the stimulus interval compared to that during the 2 s before the stimulus. (F) Response of the left and right AF7 to 0° prey stimulus. Vertical dashed lines indicate the stimulus interval. Solid line and shaded region represent the mean and 95% confidence interval, respectively (n = 7 fish). (G) Spatial tuning curves showing maximum activity of AF7 (within the 3 s time window after stimulus onset) in response to prey stimuli at different azimuths. Solid lines and shaded regions represent the mean and 95% confidence interval, respectively (n = 7 fish). Also see Figure S1.

We also found that blocking one eye affected the range of azimuths that would trigger hunting, and the kinematics of the prey capture swims. In the monocular condition, larvae would respond to prey on the exposed eye side, and up to 10° on the blocked side, but not to more lateral stimuli on the blocked side, indicating that the binocular field in our eyes-fixed condition is +/− 10° (Figure S1A). For central prey stimuli, the occlusion also caused larvae to bias the initial hunting tail beat to the exposed eye side (Figure S1B), and bout asymmetry (a measure of “J-ness”, or time the curvature of the distal part of the tail matched the overall bout direction) was higher in the monocular conditions (Figure S1C), indicating that the presence of the blocker affected hunting behavior.

### One population of prey responsive neurons is binocular

We next used two-photon imaging to identify prey-responsive neurons (PRNs) in binocular and monocular conditions. We used an electrically tunable lens to image a 500×500×100 µm volume which included the optic tectum (OT), pretectum (Pt), tegmentum (Teg), nucleus isthmi (NI), and thalamus (Thal) (Figure 2A). All these areas have previously been shown to respond to prey stimuli^22–27^. For larvae expressing nuclear-localized GCaMP6s pan-neuronally (*Tg(elavl3:Hsa.H2B-GCaMP6s)jf5Tg,* aka *elavl3*)^28^, we presented the same prey stimuli as in Figure 1 while recording calcium responses in 9 planes of the brain at 3 volumes per second, and mapped the positions of the cell bodies onto the mapZebrain atlas^29^ (Figure S2A). We moved the eye-blocker to different positions with respect to the larva to record the response to each stimulus in the left-eyed, right-eyed, binocular, and fully blocked conditions (Figures 2B and C). We also recorded eye and tail movements, and used eye convergence as a marker for hunting. Although the eyes were in agarose, we could still observe convergences of a few degrees during presentation of prey stimuli, and we used these small convergences (4.5° binocularly, on average) to classify hunting behavior (Figure 2C, Movie S1, methods). These convergence events occurred almost exclusively during prey presentation (Figure S2B). To identify PRNs, we modeled the activity of each neuron as a weighted sum of stimulus regressors (Figure S2C). We found that neurons with R^2^ ≥ 0.3 were well correlated with prey stimuli (Figure S2D), and that the fully blocked condition resulted in virtually no PRNs above this threshold (Figure S2E). Therefore, we defined neurons with R^2^ ≥ 0.3 as prey responsive. We then used spatial p-value analysis^30^ to remove neurons that were not colocalized across the 15 fish. We found that, while the majority of the responsive neurons were in the hemisphere contralateral to the unblocked eye, we also observed a smaller population of PRNs in the hemisphere ipsilateral to the unblocked eye, and we termed these neurons “ipsi-neurons” (Figure 2E).

**Figure 2.**
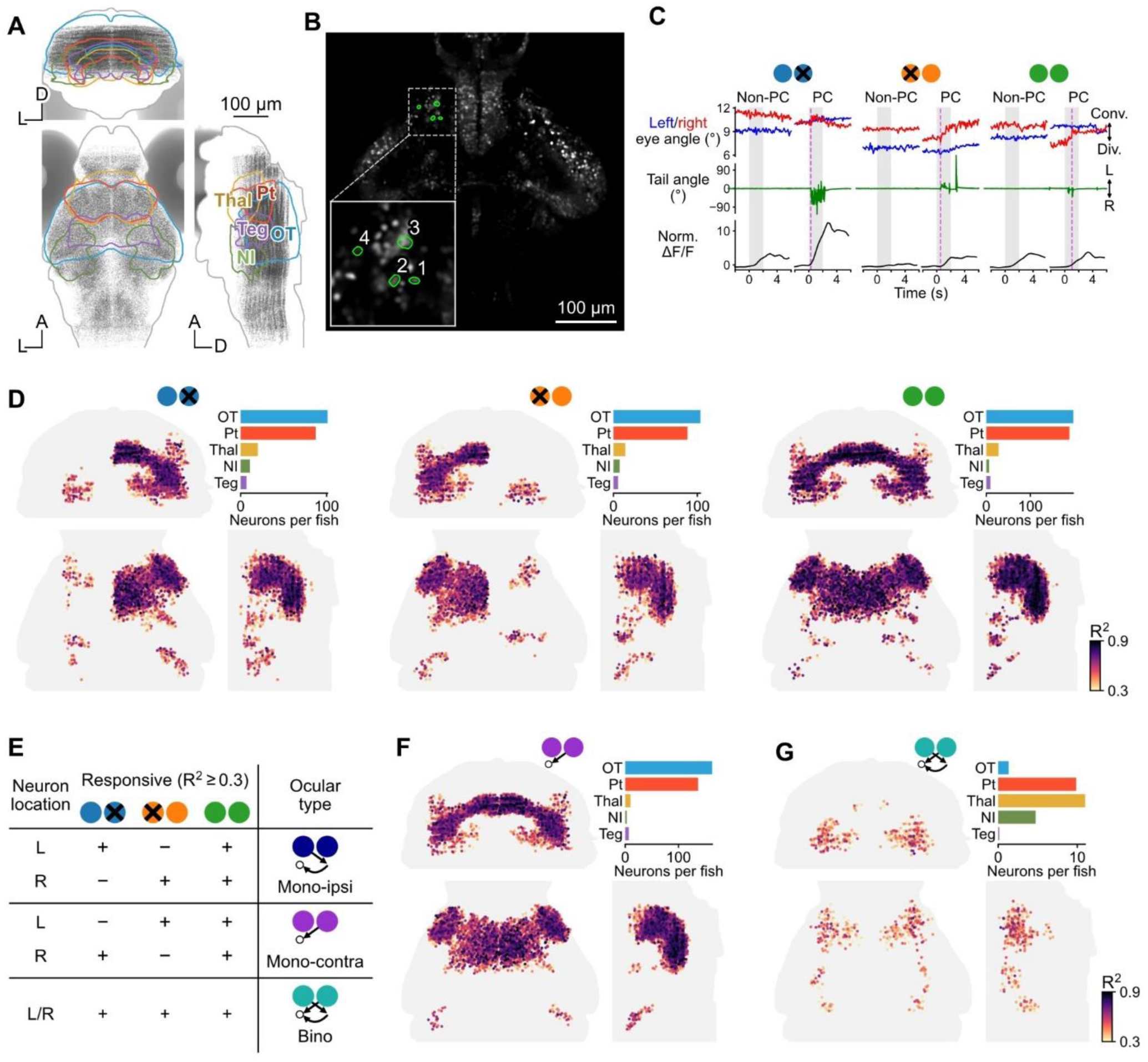
Volumetric two photon imaging in different ocular conditions identifies binocular prey-responsive neurons (PRNs). (A) Map of the cell body locations of all neurons in our imaging dataset of 15 larvae. Each grey dot represents one neuron. Coronal, horizontal and sagittal views (all anatomical maps follow the same layout unless otherwise stated). (B) Example PRNs in Pt. (C) Eye and tail movements triggered by prey stimuli and normalized ΔF/F responses of example neuron no. 1. Magenta dashed lines indicate hunting onsets (defined by eye convergence). (D) Maps of PRNs in different ocular conditions from coronal, horizontal, and sagittal views (n = 15 fish). Neurons with prey-related activity were identified with regression analysis for each ocular condition, using an R^2^ threshold of 0.3. Color code represents R^2^ value. Bar plots show average number of neurons in selected brain regions. (E) Schematic of the assignment of ocular type based on responsiveness in different ocular conditions. (F-G) Maps of the anatomical locations of monocular contra PRNs (mono-contra) and binocular PRNs (bino-PRNs). Also see Figure S2 and S3.

We then asked how the population of PRNs responded across different ocular conditions; for example, whether a particular PRN could respond to input from either eye (Figure 2E). As expected, a large class of PRNs only responded to contralateral eye input, and failed to respond to ipsilateral input (mono-contra, Figure 2F). We also found that a small population within the ipsi-neurons was unresponsive to contralateral eye input (mono-ipsi, Figure S2G). Finally, one population of ipsi neurons also responded to input from the contralateral eye, meaning that either eye could activate them (binocular, Figure 2G). We identified the anatomical locations of the binocular PRNs (bino-PRNs), and found that they were predominantly mapped to three areas: the Pt, Thal, and NI (Figure 2G, bar plot). The number and locations of bino-PRNs varied somewhat between individual larvae, but out of the 15 larvae in our dataset, 11 had bino-PRNs in the Pt and Thal, and 6 had bino-PRNs in the NI (Figure S2G).

To confirm that the bino-PRNs are not an artifact of incomplete eye occlusion, we also performed lensectomies to remove one lens of the larvae, which renders larvae unable to respond to prey^12^ (Figure S3A). The PRN patterns of sham, unilateral and bilateral delensed fish resembled those produced by the eye-blocker, and ipsi-neurons were found in the Pt, Thal and NI of unilaterally delensed larvae (Figures S3B and S3C).

### Bino-PRNs respond specifically to prey-like stimuli

Next, we investigated whether bino-PRNs respond specifically to prey, or are activated by other behaviorally relevant visual stimuli. As zebrafish primarily use red cones to detect global motion^31,32^, we introduced a red background to all visual stimuli in the section. We collected a second imaging dataset, in which larvae were first presented with circulating prey dots as in Figure 2, and then with a battery of eight additional stimuli (darkening/brightening, sweeping UV dots, horizontally moving gratings, and looming disks) in the binocular condition (Figure 3A).

**Figure 3:**
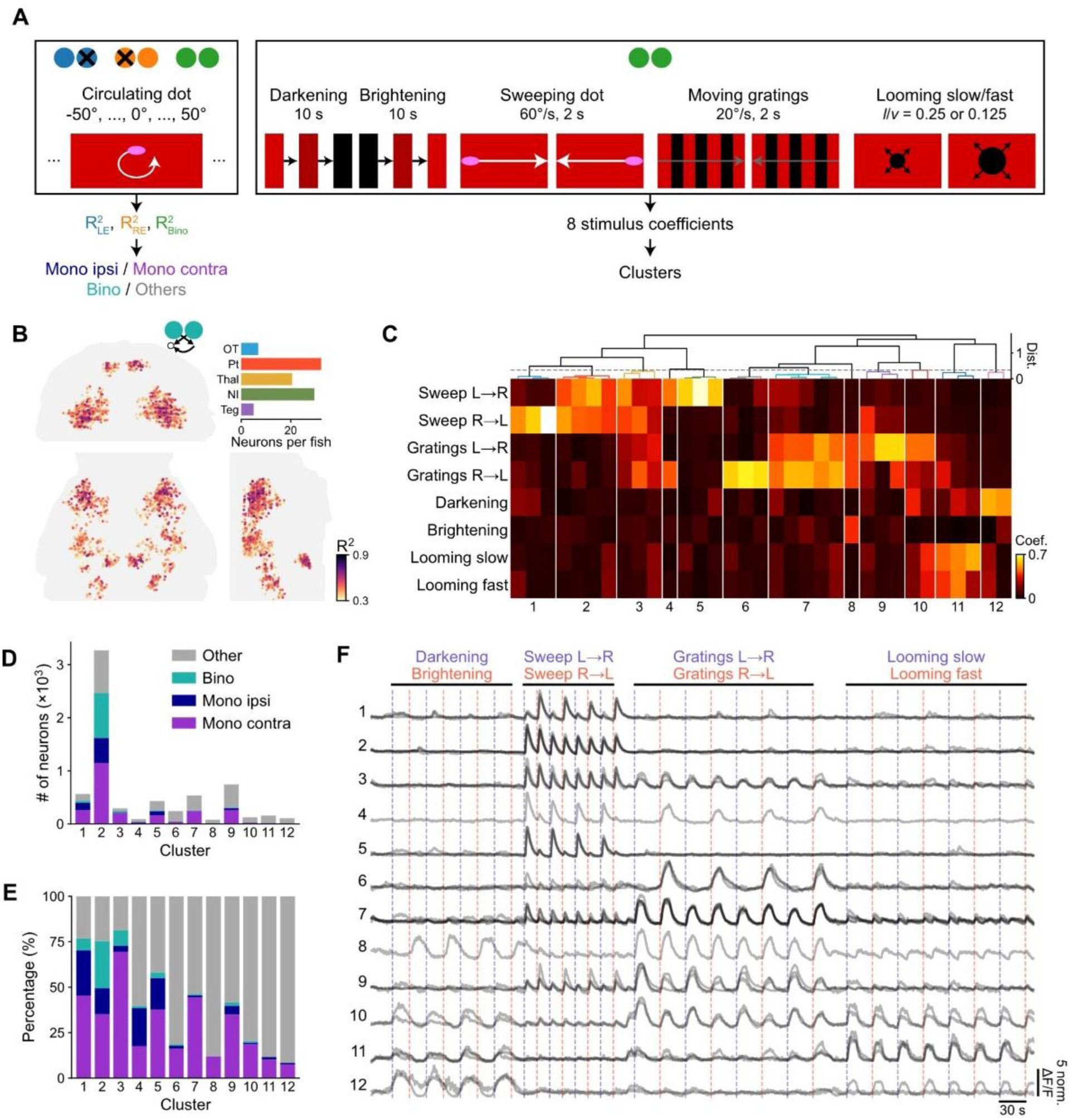
Bino-PRNs respond specifically to prey stimuli. (A) Schematic of the stimulus protocol. Circulating UV prey-like stimuli were presented on a red background in different ocular conditions, and a battery of 8 other types of stimuli were presented binocularly to assess the specificity of bino-PRNs. (B) Bino-PRNs were found in the Pt, Thal and NI using the same method as in Figure 2. (C) Hierarchical clustering revealed 12 clusters that responded primarily to prey, moving gratings, or looming stimuli. (D,E) Number and percentage of neurons in each cluster that are bino- or mono-PRNs. (F) Calcium traces of all exemplars (grey) from each cluster. (n = 10 fish)

We identified bino- and mono-PRNs as in Figure 2 (Figure 3B). Using affinity propagation clustering followed by hierarchical clustering^26^, neurons were grouped into functional types according to their responses to the eight visual stimuli (Figure 3C). The vast majority of bino-PRNs were assigned to cluster 2 (Figure 3D and E), which responded to both leftwards and rightwards moving dots (Figure 3F). In summary, bino-PRNs respond selectively to prey-like stimuli with circular motion or sweeping in both horizontal directions, rather than to moving gratings, looming stimuli, or changes in luminance.

### Bino-PRN receptive fields

To better understand the potential visual functions of the bino-PRNs, we next mapped their left eye and right eye receptive fields. Since we presented the prey stimulus at 12 different azimuths, for each neuron we can make left eye and right eye tuning curves for horizontal prey positions (Figure 4A). We find that for some neurons, the two eye’s turning curves are both close to the midline and separated only by about 10° (Figure 4A, example neuron 1), while in others, the peaks of the tuning curves were more lateral (Figure 4A, example neuron 2). We plotted all peak separations for all of our bino-PRNs, and found that there is a broad range of left to right eye peak separation distances, from 10-50° (Figure 4B and S4A). This range of peak separations could reflect neurons that are activated by objects at different distances from the hunting larva. In our imaging experiments, the eyes were fixed and could only converge a few degrees, but in natural conditions, the eyes can converge further, depending on the distance to the prey^10^. A neuron with a peak separation of 10° in our eyes-fixed imaging preparation (vergence angle ∼15°) would be optimally activated by an object 0.2 mm from the larva when the eyes are fully converged (Figure 4C, blue lines), whereas a neuron with a peak separation of 50° would be optimally activated by an object 1.29 mm away (Figure 4C, violet lines). When the eyes are partially converged, these neurons would be best activated by more distant objects (Figure 4C, 60° vergence), with a range similar to the maximum detectable prey distance (Calculation: Figure S4B). We also plotted the distributions of bino-PRN peak separations for each brain area, and found that the distributions were similarly shaped, with a peak around 30° of left eye to right eye separation (Figure S4C). Thus, bino-PRNs in all five regions have left and right eye receptive fields that are offset by varying degrees, over a range corresponding to naturalistic prey distances, and could thus combine binocular visual information with the degree of convergence to determine prey position.

**Figure 4.**
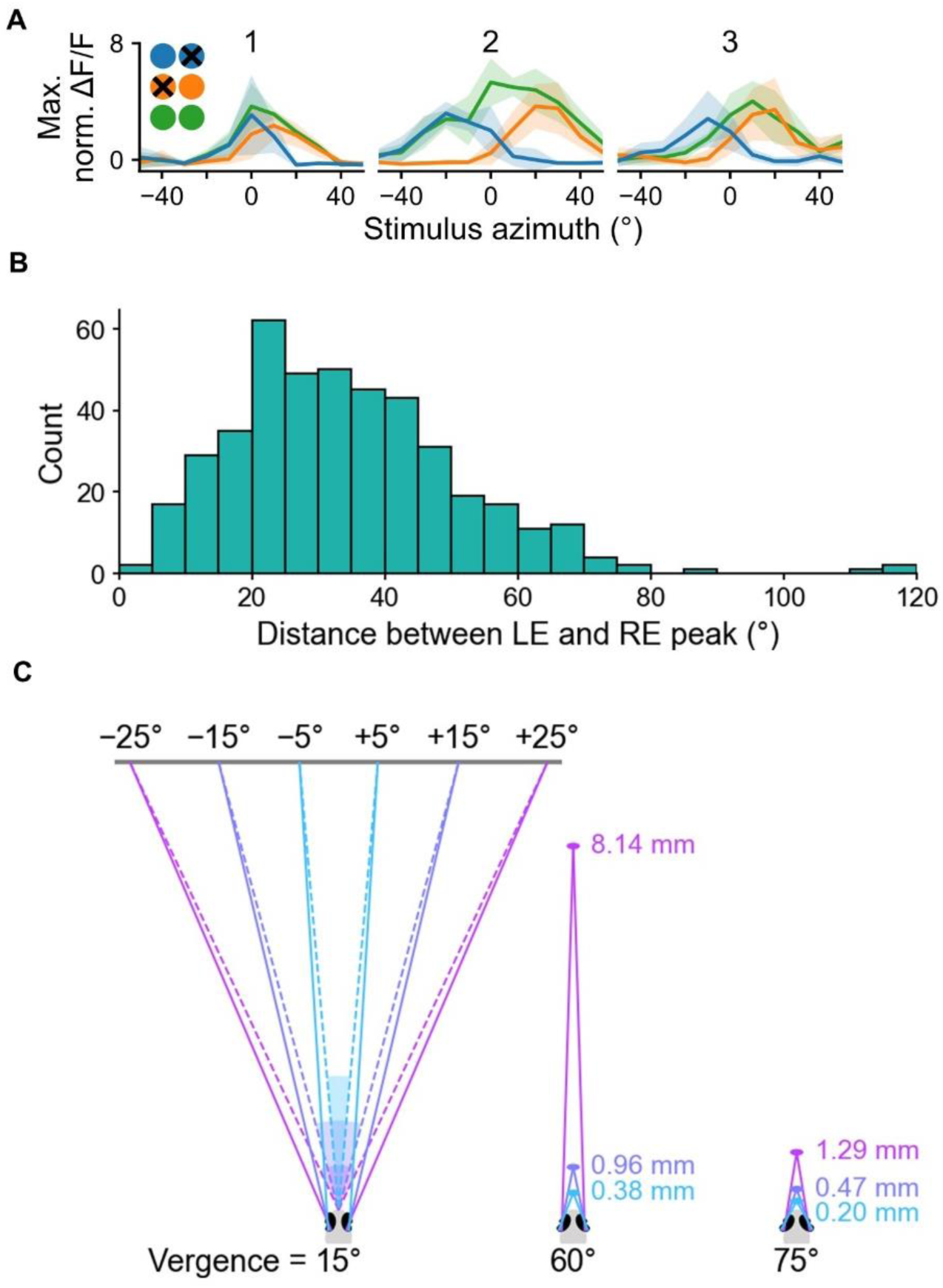
Bino-PRNs receive inputs from the two eyes which are offset in horizontal space, and correspond to objects at different distances from a hunting larva. (A) Monocular and binocular tuning curves of three example bino-PRNs from one larva. Blue line represents the left eye response to stimuli presented at each azimuth, orange line represents the right eye response, and green is the binocular condition. Shading represents 1 SD. (B) Distribution of the distances between the peaks of the left eye and right eye receptive fields for all bino-PRNs. (C) Schematic showing how left eye/right eye peak separations in the imaging preparation correspond to object distances at medium (60°) and maximum (75°) levels of convergence. Blue lines = 10° separation, purple lines = 30° separation, violet lines = 50° separation between right and left eye peaks. Also see Figure S4.

### Bino-PRNs show enhanced activity during hunting trials

Bino-PRNs appeared in some cases to respond more robustly in hunting trials than non-hunting trials (Figure 2C), and to further investigate this property, we plotted the responses of bino-PRNs in each brain region (Figure 5A) in prey capture (PC) vs. non-prey capture (non-PC) trials. Indeed, the bino-PRNs in each area were on average more activated in prey capture trials (red lines, Figure 5B). Thus, one possibility could be that these neurons are binocular as a result of receiving a motor signal, rather than visual input from both eyes. However, their binocularity was not entirely a result of the hunting state, as they were activated by prey stimuli seen by the ipsilateral eye even in non-hunting trials (Figure 5B, middle column, black lines). In contrast, the contra-PRNs in the OT, which constitute the majority of the PRN population, had no response to ipsilateral prey, even in hunting trials (Figure 5B, lowest row). This suggests that the bino-PRNs are receiving sensory input from both eyes, as well as responding more strongly in hunting trials.

**Figure 5.**
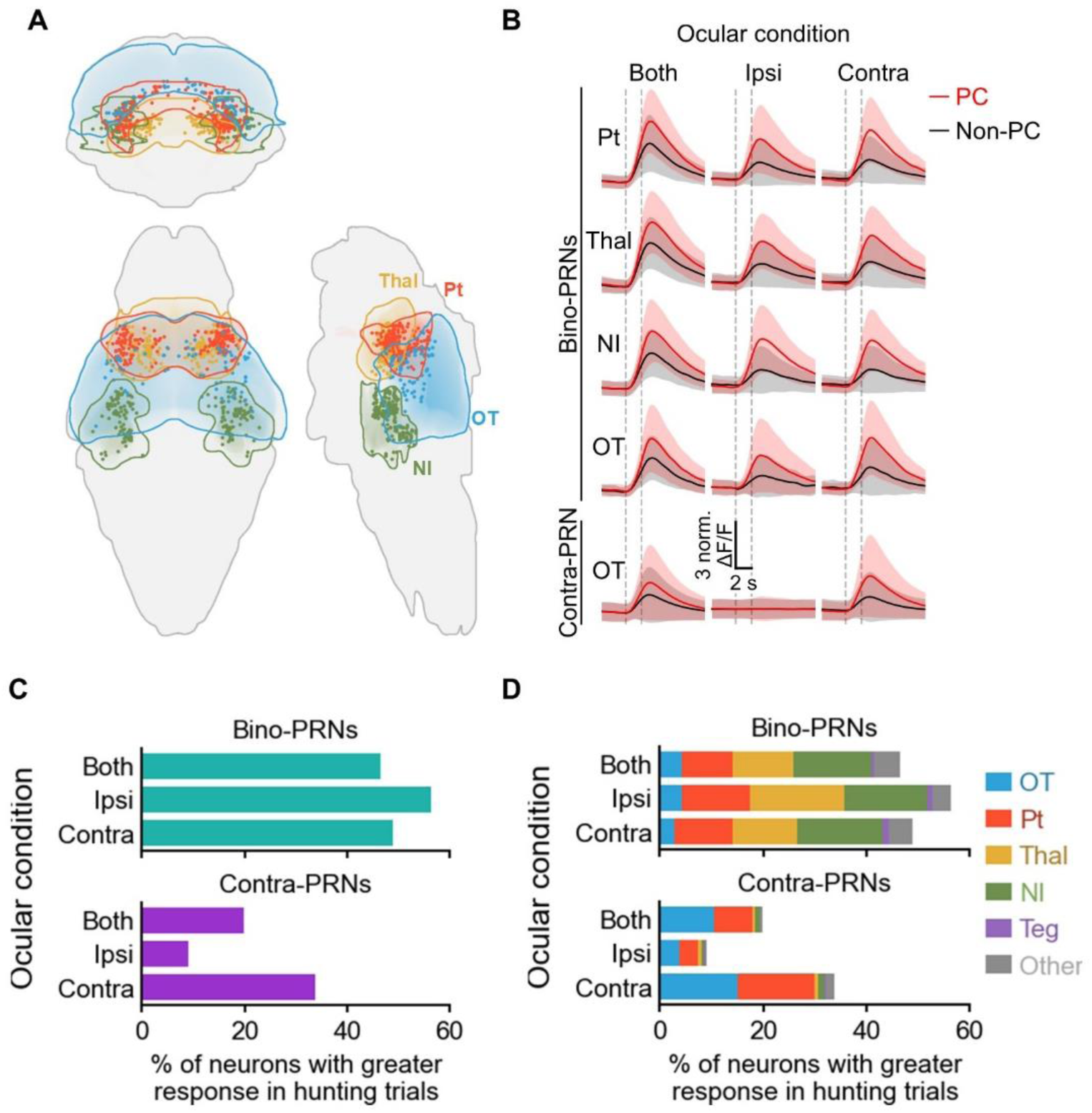
Bino-PRNs are motor modulated and receive binocular input in no behavior trials. (A) Anatomical locations of bino-PRNs in four selected regions of the brain; coronal, horizontal, and sagittal views. (B) Average responses of bino-PRNs in each region and mono-contra neurons in the OT to prey capture (PC) and non prey capture (non-PC) trials (defined by eye convergence) to prey at 0° in different ocular conditions. In the Ipsi condition, only the ipsilateral eye relative to the PRN was able to see the stimulus. Solid lines and shaded regions represent mean and ±1 SD, respectively. (C) Percent of bino- and contra-PRNs with significantly larger responses during hunting trials. Trial response is calculated as the mean dF/F between +1 and +3 seconds after stimulusa onset. A neuron’s response is considered significantly greater if the one-sided Mann-Whitney U test comparing hunting trial responses to NR trial responses results in a p-value less than 0.05. (D) Proportion of bino- and contra-PRNs with significantly greater responses in hunting trials in each brain area.

We next looked at motor enhancement at the individual neuronal level by asking whether individual PRNs respond significantly more in hunting than non-hunting trials. We find that a large fraction of bino-PRNs are significantly more responsive during trials with eye convergence (48% in the binocular condition), whereas a smaller percentage of contra-PRNs (20% in the binocular condition) show a significant enhancement (Figure 5C). This indicates that the motor enhancement is not a global signal to all neurons or even all PRNs, but is more robust in the bino-PRN population. We looked at the locations of these motor-modulated bino-PRNs, and found that they were not confined to one brain area, but distributed among the Pt, Thal, and NI, the three areas with the largest number of bino-PRNs (Figure 5D). The finding that bino-PRNs are more responsive during hunting suggests that they are either generating the hunting motor command, or receiving a corollary discharge when hunting is initiated.

### The *lhx9:Gal4* line labels bino-PRNs

We next sought to identify Gal4 lines labeling the bino-PRNs to allow us to use a genetic marker to image and manipulate these neurons. We therefore performed functional imaging of several lines in different ocular conditions. We first tested the *KalTA4u508* line, which was reported to express KalTA (a Gal4 variant) in the hunting command-like neurons of the accessory pretectal nucleus (APN) (Figure 6A)^23^. It is possible that these command-like neurons could be integrating input from the two eyes. However, when we imaged *Tg(KalTA4u508;UAS:GCaMP6s)* larvae, we found that almost all PRNs were responsive only to the contralateral eye prey stimulation, and were classified as mono-ipsi or mono-contra (Figure 6B-D). All of the pretectal PRNs in the line were mono-contra, meaning that the pretectal command-like neurons must be receiving input from only the contralateral eye.

**Figure 6.**
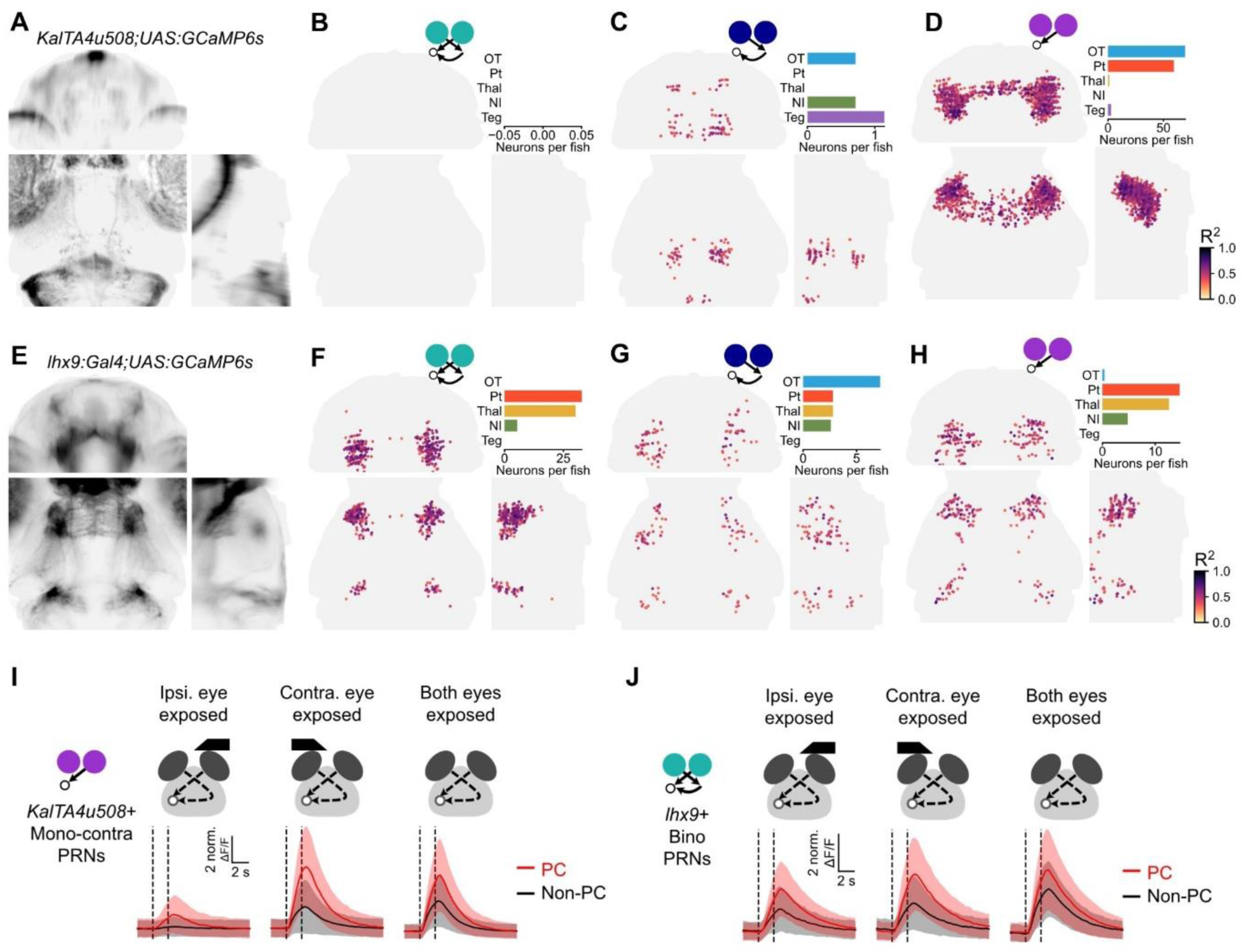
Identification of a transgenic line labelling bino-PRNs. (A) Expression pattern of the KalTAu508 line. (B,C,D) Binocular, mono-ipsi and mono-contra PRNs in the u508 line (n = 7 fish). (E) Expression pattern of the lhx9 line. (F,G,H) Binocular, mono-ipsi and mono-contra PRNs in the lhx9 line (n = 5 fish). (I) Average response of the mono-contra PRNs in the KalTA4u508 line in prey capture (PC) and non prey capture (non-PC) trials. Solid lines and shaded regions represent mean and ±1 SD, respectively. (J) Average response of the bino-PRNs in the lhx9 line. Solid lines and shaded regions represent mean and ±1 SD, respectively.

In contrast, when we imaged *Tg(lhx9:Gal4;UAS:GCaMP6s)* larvae^33^ (Figure 6E), we found that the majority of PRNs were binocular. The *lhx9+* bino-PRNs were located in the Pt (as well as Thal) and the anterior parts of the NI (Figure 6F-H). These neurons were activated by prey presented to either the contra- or ipsilateral eye (Figure 6J). Indeed, these neurons were activated by ipsilateral prey even in the absence of hunting, similar to the bino-PRNs in the panneuronal dataset (Figure 5B). Taken together, these data suggest that the *lhx9+* bino-PRNs represent a separate population that is not labelled by the *KalTA4u508* line. We can therefore use these two lines to investigate the source of the motor enhancement of the bino-PRNs.

### Optogenetically triggered hunting events activate *lhx9*+ Pt, Thal and NI neurons in the absence of prey

Based on the enhanced activity of bino-PRNs in hunting trials (Figure 5), we hypothesized that they could be activated as a consequence of hunting. Zebrafish larvae normally have unconverged eyes, to maximize coverage of the visual field, and only converge their eyes at the onset of a hunting episode^9,10,13^. It may be that in order for the brain to properly interpret binocular visual input, some information about hunting onset or the degree of convergence has to be conveyed to visual areas. One way for this information to get there would be via corollary discharge. discharge, or an internal signal that is the result of a motor command^34^. In this case, visual areas could receive a signal that is the result of the hunting motor command, that would indicate that the eyes are converged, and the animal is hunting. However, it is also possible that PRNs are more responsive during hunting trials because they are playing a role in generating the motor command.

To distinguish between these possibilities, we used holographic optogenetic activation of hunting command-like neurons to artificially induce prey capture (opto-hunting), and then recorded the activity of PRNs to observe whether they receive an excitatory input in opto-hunting trials. The *KalTA4u508* line was previously shown to label neurons in the APN whose activation can evoke prey capture^23^. We expressed GCaMP6s and Channelrhodopsin (ChR2) in the *u508* and *lhx9+* neurons (*Tg(KalTA4u508;lhx9Gal4;UAS-GCaMP6s;UAS:ChR2(H134R)-mCherry)* larvae). We used 920 nm 2-photon 3D-holographic stimulation to activate a small area (12 x 12 µm, Figure S5A-D) within the APN of these larvae, which resulted in a calcium response in these neurons (Figure S5E and F). At the same time, we imaged GCaMP at 1020 nm (Figure 7B, left)^35^. The optogenetic sessions were followed by a visual stimulation session, in which the fish was presented with prey-like stimuli (Figure 7B, right). We used eye convergence to define hunting events, and found that these events occurred during the optogenetic experiments, both in and out of phase with the holographic stimulation (Figures 7C, middle panel, red dashed lines, and S6A). These bouts were characterized by eye convergence and J-turn or forward swim tail movements (Figure S6B).

**Figure 7.**
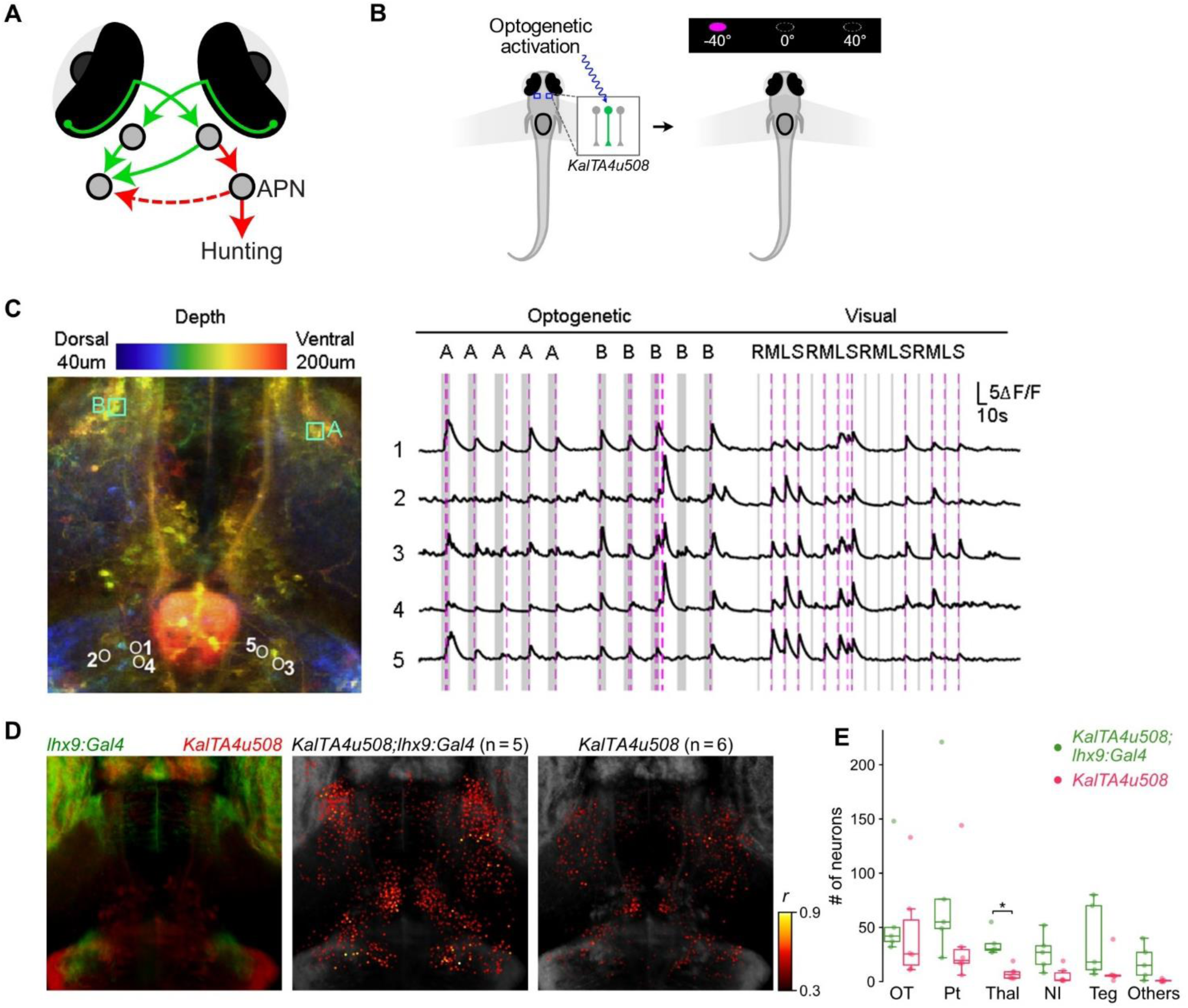
Optogenetically triggered hunting events activate PRNs. (A) Model of motor signal from command-like APN neurons to bino-PRNs. (B) Schematic of sequential optogenetic activation and visual stimulation paradigm. KalTA4u508+ Pt neurons were holographically activated to trigger prey capture while performing 2-photon calcium imaging. After the opto-imaging session, prey stimuli were presented to record visual responses. (C) Left: example larva expressing GCaMP6s and ChR2 under control of the KalTA4u508 and lhx9 lines. Locations of the stimulation volumes (cyan squares, 12 x 12 µm) and opto-hunting correlated neurons (circles). Middle: calcium traces of the top 5 opto-hunting correlated neurons in one fish. Grey shading represents stimulation of the right (A) or left (B) APN. Magenta dashed lines represent hunting onsets. Right: Responses to visual prey stimuli in the same 5 neurons to right, middle, and left circulating prey and sweep prey stimuli (RMLS). (D) Left: the lhx9 (green) and u508 (red) expression patterns. Locations of opto-hunting neurons in larvae with both gal4 lines (Middle) vs. the u508 line only (Right). (E) Numbers of opto-hunting neurons in larvae with both gal4 lines vs. the u508 line only (Mann-Whitney U tests with Bonferroni correction, *P < 0.05, otherwise P > 0.05). Also see Figure S5, S6 and S7.

To determine whether these hunting bouts were due to the two-photon scanning activating ChR2, we performed control experiments in larvae with different combinations of the three transgenes. We found that the ChR2 and *KalTA4u508* transgenes were essential for the induction of hunting (Figure S6C). In fish carrying these two transgenes, we did observe hunting bouts in experiments with scanning but no stimulation (Figure S6C, orange bar), suggesting that these bouts were due to activation of ChR2 by the imaging laser. Indeed, in the absence of two-photon scanning and holographic stimulation, we observed no spontaneous prey capture bouts (Figure S6C, green). In summary, our holographic stimulation can evoke hunting, but two-photon scanning can also activate the *KalTA4u508* APN neurons to cause some prey capture bouts during the scanning period.

We observed that when our activation of the command-like neurons induced hunting, certain GCaMP-expressing cell bodies were activated. To identify neurons correlated with opto-hunting behavior, we constructed regressors for prey capture events, and defined neurons with R^2^ ≥ 0.3 as opto-hunting neurons. After activation of a small area within the APN (Figure 7C, left panel, squares), we found cell bodies that were highly correlated with optogenetically-evoked eye convergence (Figure 7C, middle panel). Many of these neurons also responded to our visual prey stimuli (Figure 7C, right panel), indicating that they are PRNs. We also performed the experiment in larvae with only the *KalTA4u508* Gal4 driving ChR2 and GCaMP6s expression, and compared them to larvae with both Gal4s (Figure 7D, right). We found that larvae with both Gal4s had more opto-hunting neurons in the Thal, Pt and NI (Figure 7E, green) than those with *u508* only (red). This suggests that the *lhx9* neurons are downstream of the hunting motor command. We also asked whether the opto-hunting neurons responded to visual stimuli, and found that in the thalamus, pretectum and NI, about one third of the opto-hunting neurons were also classified as PRNs (Figure S6D). To determine how many of the opto-hunting neurons identified in this experiment were *lhx9+*, we created masks of the brain regions labeled by the two lines (Figure S6E, left, methods), and found that many of the opto-hunting neurons were located within the *lhx9* mask (Figure S6E, middle), and these neurons were mostly located in the Thal, Pt, and NI (Figure S6F). These results suggest that the lhx9-positive PRNs are activated by the hunting command, possibly via corollary discharge.

One caveat of the *u508/lhx9* experiments is that it is possible that some opto-hunting bouts are triggered by the scan, and although the *u508* line was required for opto-hunting (Figure S6C), we did not completely separate the stimulated and responding neuronal populations. To address this concern, we performed similar optogenetic experiments with GCaMP expressed in a separate population of neurons from the *u508* line. We used larvae with GCaMP expressed panneuronally, and ChR2 only in the *u508* line (*elavl3:Hsa.H2B-GCaMP6s* and *KalTA4u508;UAS:ChR2(H134R)-mCherry*), and repeated experiment in Figure 6B with these fish. In this larger population, we found that the opto-hunting neurons shared a similar anatomical pattern with the bino-PRNs identified in Figure 2H (Figure S6G, left). In addition, a subset of these neurons were located within the *lhx9* mask (Figures S6G, right), and approximately half of these opto-hunting neurons responded to visual prey stimuli and were classified as PRNs using the same threshold as for Figure 2 (Figure S6H).

Due to space constraints in the holographic optogenetic setup, we were not able to use the eyeblocker to directly determine how many of the opto-hunting neurons were bino-PRNs. However, we show that PRNs in the Thal, PT and NI are activated as a consequence of the hunting motor command, and many of these neurons were within the *lhx9* line, which labels substantially more bino-than contra- or ipsi- PRNs (Figure 2F-H). This suggests that the motor modulation of the bino-PRNs (Figure 5B-C) is likely due to a hunting-related corollary discharge.

### Coactivation of command-like neurons and lhx9 evokes prey capture swims

We next sought to identify the functional role of the lhx9+ neurons by optogenetically activating them and observing the behavioral outcome. We first used an optic fibre to deliver 470 nm light to the whole brain to activate *KalTA4u508* neurons (Figure S7A). As was previously reported, activating *KalTA4u508* neurons drove larvae to perform prey capture J-turns and forward swims^23^. We then did the same optic fibre activation in larvae with Channelrhodopsin (ChR2) expressed in both the *lhx9* and *KalTA4u508* populations, which produced hunting bouts (classified by eye convergence) that resembled turns (asymmetrical hunting) or forward swims (symmetrical hunting), as well as non-hunting bouts (other, Figure S7B, Supplementary Video 2). In the double Gal4 larvae, we found that the optogenetically evoked prey capture bouts were significantly more symmetrical than for *u508* only (Figure S7C). Larvae bearing only the *lhx9* Gal4 also had higher bout symmetry (Figure S7B). This higher symmetry could be due to a larger proportion of forward swim bouts, or a higher degree of symmetry within the forward swim bouts. However, a caveat of this experiment is that although PRN population within the *lhx9* line consists predominantly of bino-PRNs (Figure 6), the line also contains non-PRNs, and so we cannot ascribe the symmetrical prey capture bouts to bino-PRN activation. We then performed two-photon ablations of lhx9+ PRNs to determine whether they are mediating the change in swim kinematics. As the lhx9+ PRNs in the thalamus and pretectum were difficult to ablate without damaging the eyes, we ablated ∼20 lhx9+ PRNs in the NI of double Gal4 larvae and then performed the optic fibre experiment. We found that the ablated larvae had more asymmetrical swims than the sham controls (Figure S7D), suggesting that the lhx9+ PRNs, which are largely bino-PRNs, contribute to the more symmetrical opto-hunting bouts of the double Gal4 fish compared to *u508* only. One possibility is that bino-PRNs are involved in the approach phase of prey capture, which typically occurs after the initial turn, when the prey is visible to both eyes. However, a more precise assignment of the behavioral role of bino-PRNs will require selective activation of these neurons.

## Discussion

Here, we identify binocular prey-responsive neurons in the zebrafish visual system by sequentially occluding the eyes and volumetrically imaging visual brain areas. We classified cell bodies that responded to prey stimuli as PRNs, and those that responded when the stimulus was presented to the left and the right eye (separately) as bino-PRNs. We found that these neurons are located primarily in the Pt, Thal, and NI. Notably, our bino-PRN population contained very few neurons in the optic tectum (Figure 2H), although the tectum had the largest share of mono-contra PRNs (Figure 2G). This absence is somewhat surprising given the importance of the tectum in prey capture behavior^22^. It may be that while the tectum is important for some aspects of hunting, the Pt, Thal, and NI are the loci of excitatory binocular interactions. However, the tectum may receive inhibitory interhemispheric input, as shown by a recent study that identified a population of intertectal neurons that provide inhibitory input bilaterally to the OT^25^.

The identification of binocular neurons allows us to analyse their functional properties to evaluate their potential roles in prey capture. We found that the left eye and right eye receptive fields of the bino-PRNs were offset to varying degrees, with the peak of the distribution at an offset of 30° of the visual field (Figure 3B). In a freely swimming larva, the eyes would converge during hunting, and in the case of full convergence, the bino-PRNs with a 30° offset would be best activated by prey objects 0.47 mm from the larva. Larvae generally converge their eyes fully near the end of a hunting episode, when they have approached the prey and placed it in the central visual field, and they then typically execute a strike when the prey is 0.5 mm away^12^. The fact that bino-PRNs are tuned for prey objects at the naturalistic strike distance suggests that they could allow the larva to estimate distance to the prey in order to successfully capture it.

Our imaging data also suggested that bino-PRNs were more active in trials where hunting was initiated (Figure 4). This effect could occur because the bino-PRNs are involved in generating the hunting motor command, or because they are receiving the hunting motor signal. We hypothesized that PRNs could receive a corollary discharge as a result of hunting initiation, and to test this hypothesis, we asked whether artificial induction of hunting in the absence of visual stimuli would also activate PRNs. Indeed, in trials where hunting was optogenetically triggered, a number of neurons in the Pt, Thal, and NI were activated, and many of these neurons were PRNs (Figure 6). Due to technical limitations, we were not able to use the eyeblocker to identify bino-PRNs in the holographic optogenetic setup, so cannot directly say how many of the opto-hunting neurons were bino-PRNs. However, our data indicate that bino-PRNs are more likely than contra-PRNs to be motor enhanced (Figure 4C), and that PRNs in the three areas containing large numbers of bino-PRNs are activated by the hunting motor command, making it likely that the bino-PRNs are receiving a hunting-related motor signal.

As the signature of hunting is eye convergence, the motor signal could indicate the degree of convergence, which would be informative for estimating distance to prey. If larvae are using triangulation, where activation of different populations of bino-PRNs corresponds to prey at different distances, it would also be crucial to know how converged the eyes are, as the same set of neurons would be activated by more distant prey when the eyes are not fully converged (Figure 3C, 60° binocular angle). However, convergence angle information would also be informative for disparity-based distance estimation, as it would indicate the distance of the fixation point, which must be combined with disparity information to determine distance to the prey^15,36^. Therefore, further experiments will be required to elucidate the nature of the binocular computation.

Overall, we have identified for the first time a set of binocular prey-responsive neurons which zebrafish larvae could use to successfully capture prey. Based on their receptive fields, the majority of these neurons respond optimally to prey at striking distance from a hunting larva, and they may receive a corollary discharge signal indicating eye convergence. These properties could allow the larva to determine whether to execute a strike, and with what degree of propulsion, or whether an additional approach swim is required to reach strike range. Additional experiments will be required to determine the exact role of binocular neurons in prey capture, how these neurons receive input from the ipsilateral eye, and the circuitry and significance of the motor signal, and the type of binocular computation being implemented. However, our findings lay the foundation for understanding binocular vision in a simple vertebrate visual system.

## Materials and Methods

### Animals

All procedures conformed to the guidelines by the animal ethics committee of HKUST. Animals were kept under a standard 14:10 light cycle at 28°C. Zebrafish larvae carrying mutations in the *mitfa* allele (nacre) were used for all experiments^37^. Larvae were kept in Danieau’s solution, and fed with paramecia (*Paramecium multimicronucleatum*, Carolina Biological Supply Company) on 5 to 7 days post fertilization (dpf).

The following transgenic lines were used for this study: *Tg(atoh7:GAL4-VP16)s1992tTg*^38^*, Tg(UAS:GCaMP6s*)*mpn101*^39^*, Tg(elavl3:Hsa.H2b-GCaMP6s)jf5Tg* ^28^*, Tg(KalTA4u508)u508Tg*^23^*, Tg(lhx9:Gal4VP16)mpn203*^33^*, Tg(UAS:ChR2(H134R)-mCherry)mpn134*^35^. For imaging of RGC axons and pan-neuronal imaging, For imaging of RGC axons and pan-neuronal imaging, *Tg(atoh7:Gal4VP16;UAS:GCaMP6s)* and *Tg(elavl3:Hsa.H2B-GCaMP6s)* fish were used, respectively*. Tg(lhx9:Gal4VP16;UAS:GCaMP6s)* and *Tg(KalTA4u508;UAS:GCaMP6s)* larvae were screened for binocular neurons. For two-photon optogenetic stimulation with calcium imaging experiments, *Tg(KalTA4u508;lhx9:Gal4VP16;UAS:GCaMP6s;UAS:ChR2(H134R)-mCherry), Tg(KalTA4u508;UAS:GCaMP6s;UAS:ChR2(H134R)-mCherry), Tg(lhx9:Gal4VP16;UAS:GCaMP6s;UAS:ChR2(H134R)-mCherry) and Tg(elavl3:Hsa.H2B-GCaMP6s; KalTA4u508;UAS:ChR2(H134R)-mCherry)* fish were used. For optic fiber stimulation experiments*, Tg(KalTA4u508;lhx9:Gal4VP16;UAS:ChR2*(*H134R)-mCherry), Tg(KalTA4u508; UAS:ChR2(H134R)-mCherry)* and *Tg(lhx9:Gal4VP16;UAS:ChR2(H134R)-mCherry)* fish were used.

### Zebrafish Preparation

6-8 dpf larvae were embedded in 2% low melt agarose (Invitrogen) on 100 mm square (Thermo Scientific (cat# 109-17)) or 90 mm round (Thermo Scientific (cat#101VR20)) plastic petri dish. For eye-blocker experiments, the agarose surrounding the tail and in front of the head was removed by surgical blade (Paragon p303) to allow for tail and fin movements while eyes were kept in the agarose. Otherwise, both eyes and tail were freed. Fish were allowed to recover overnight prior to the experiment.

### Eye-blocker design

Files describing the design of the eye-blocker are deposited in https://github.com/SemmelhackLab/eye-blocker-model. The components of the eye-blocker were fabricated using a 3D printer with black PLA plastic as the material. The eye-blocker was assembled from two arm parts and a body part. During imaging experiments, the tips of the eye-blocker arms were placed between the objective and the bottom of the fish arena. The height of the tips was chosen to minimize the gaps at the top and the bottom without hindering the vertical movements of the objective. The body of the eye-blocker was attached to a 3-axes stage via an optical post. By controlling the movements of the 3-axes stage along the left-right axis of the fish, the arms of the eye-blocker could be moved to the position required for each ocular condition. The movements of the eye-blocker were visually guided by the live preview from the high-speed camera.

### Lensectomy

At 6-7 dpf, larvae were embedded in 2% low melting agarose, and then the prep was immersed in 0.02% tricaine (MS-222 Sigma-Aldrich) to anaesthetize the larvae throughout the surgery. Lensectomies were performed with needle (Terumo 0.40 x 13mm). An incision was made in the transparent cuticle of the temporal side of the eye and lens was scouped out gently. Animals were then released from the agarose, allowed to recover overnight, and fed paramecia. The delensed larvae were re-embedded the next day (7-8 dpf), and tested the day after (8-9dpf).

### Visual Stimulation

We adopted the spatial visual stimulator module (DLP® Light Crafter™ 4500 modified by EKB Technologies Ltd) to present stimuli as previously described^40^. 385 nm “UV” LED (SOLIS-385C Thorlabs) and 590 nm “Red” LED (M590L4 Thorlabs) were used as the light sources. The Light Crafter (LCr) back projected to the 50 mm wide x 5 mm height Teflon fabric screen at 13.5 cm away. The screen was kept at 1 cm in front of the fish head.

### Design of Visual Stimuli

Visual stimuli were created using PsychoPy. The prey-like circulating stimuli were UV dots, sized 4°×2°, moving in an elliptical path of 8°×4° at a speed of 2 Hz for 2 seconds on a dark background. The long axis of the dots was aligned to the movement direction. In the eye-blocking experiment (Figures 1-4), the stimuli were presented at different positions, ranging from −50° (left) to 50° (right) in 10° steps. Stimuli on the left side moved counterclockwise, while those on the right moved clockwise. For the center azimuth (0°), we presented both counterclockwise and clockwise directions (indicated as −0° and +0°), resulting in 12 distinct stimuli. We randomly presented these 12 stimuli four times in each imaging session, with a 13 s interstimulus interval (ISI). We conducted two sessions for each eye condition, with at least a 5-minute pause between sessions, and followed a palindromic sequence for the eye conditions. The first stimulus in all experiments was shown 30 seconds after starting the two-photon scanning.

In the stimulus specificity experiment (Figure S4), we conducted one imaging session each for left-eyed, right-eyed, and binocular conditions. The stimuli were the identical to those used in the eye-blocking experiment, except the background was red. After the sessions with the circulating dots, we presented a battery of eight additional types of stimuli under the binocular condition: darkening and brightening of the screen, leftwards and rightwards sweeping dots, leftwards and rightwards moving gratings, and slow and fast looming disks. The darkening and brightening stimuli changed the screen from red to black, or the reverse, over 10 seconds, with an ISI of 10 s. The sweeping dots have a size of 4°×2° and moved horizontally from −60° to 60°, or the reverse, over 2 s. The grating pattern was a sinusoid alternating between red and black with a cycle of 15° moving horizontally at 20°/s for 10 s. It remained visible but stationary during the 10-second ISI. The looming stimuli were dark circles that expanded from the screen’s center at slow (l/v ratio = 0.25) and fast (l/v ratio = 0.125) speeds, with a 30-second interval between their initiations.

In the optogenetics imaging experiment (Figure 6), we conducted a visual stimulation session after the optogenetic stimulation sessions. We used a sweeping stimulus in addition to the −40°, 0°, and 40° circulating stimuli. The sweeping stimulus was a 4°×2° UV dot moving horizontally from −60° to 60° over 2 s. The stimulus sequence of (−40°, 0°, 40°, sweep) was presented four times with an ISI of 13 s.

### Behavioral Recording

Behavior was recorded at 200 Hz using a high-speed camera (PhotonFocus MV1-D1312-160-CL) at a resolution of 544×560 pixels, covering the entire fish and a small area in front to track the eye-blocker. Illumination was provided by an 850 nm infrared LED array beneath the fish arena, and an infrared band-pass filter on the camera lens blocked light from the two-photon laser.

### Calcium Imaging

For the eye-blocker experiments, in-vivo calcium imaging was performed using a two-photon microscope (Nikon A1R MP+) equipped with a 25× water-immersion objective (Nikon CFI75 Apochromat 25XC W 1300, 1.1 numerical aperture). The excitation laser was calibrated to a wavelength of 1020 nm. An electrically tunable lens (Optotune EL-16-40-TC) was mounted behind the objective to enable multi-plane functional imaging. The imaged volume contains 9 planes with an average spacing of 15 µm between consecutive planes. Each plane was imaged at a resolution of 512×512 pixels (0.99 µm/pixel) at 3 Hz. Anatomical z-stacks were collected after the imaging sessions.

For the optogenetics experiments as in Figure 5, a home-built two-photon light scanning microscope controlled by a C# based custom software is employed. The laser source is a Ti:Sapphire femtosecond laser (Chameleon Ultra II, Coherent) operating at 1020 nm wavelength for calcium imaging to reduce the crosstalk between GCaMP6s and ChR2. The laser beam first passes through the half-wave plate to adjust polarization direction. The laser beam with 2.2 mm in diameter (propagate a large distance from the laser source and expanded from 1.2mm) is expanded by a factor of ×7.14 through a 4f system (AC254-035-B-ML F = 35 mm, Thorlabs and AC254-250-B-ML F = 250 mm, Thorlabs) to full fill the ETL (EL-16-40-TC-NIR, Optotune AG), which is used for multi-plane functional imaging. Then a 4f system (AC254-200-AB-ML F = 200 mm, Thorlabs, and AC254-050-AB-ML F = 50 mm, Thorlabs) reduces the beam diameter to 4 mm to avoid overfilling the scanner mirror. A resonant scanner (CRS 8 kHz, Cambridge technology) for high-speed scanning is conjugated with a set of XY galvo scanners (6215H, Cambridge technology) by a 1:1 relay (AC254-100-AB-ML F = 100 mm, Thorlabs). Finally, two lenses (Scan lens SL50-CLS2 F = 50 mm, Thorlabs, and Tube lens TTL200MP F = 200 mm, Thorlabs) are used to correct the aberration and expand the laser beam to fill the objective back pupil (objective lens (20x, N.A. 1.0, XLUMPLFLN20XW, Olympus Corporation)). The ETL, resonant scanner, XY galvo scanners, and objective pupil plane are all conjugated. In the fluorescence collection path, a dichroic mirror (FF705-Di01-25*36, Semrock) is set between the tube lens and objective to reflect the fluorescence light, which then passes through two relay lenses (AC254-150-AB-ML F = 150 mm, Thorlabs and AC254-040-AB-ML F = 40 mm, Thorlabs) into the PMT (H11461-03, Hamamatsu). Two filters (FF03-525/50-25, Semrock, and FF01-600/52-25, Semrock) are installed on a flipper which can switch the filter to collect different fluorescence signals. Voltage signals are converted from PMT output current using a high-speed current amplifier (DHPCA-100, Femto) and digitized using an oscilloscope device (PCIe-8512H, ART technology).

### Two-photon optogenetic stimulation

The laser source is a high-power ultrafast fiber laser (Toptica & FF Ultra 920). The laser beam first passes through the half-wave plate to adjust polarization direction. And then pass through a set of galvo scanner (6215H, Cambridge Technology). Then the light beam is expended by two lenes (AC254-50-AB-ML F = 50 mm, Thorlabs, and AC254-250-AB-ML F = 250 mm, Thorlabs) to fill the spatial light modulator (HSP1K-500-1200-PC8, Meadowlark Optics). Then the SLM is conjugated to the back focal plane of objective by a demagnify relay (AC508-250-AB-ML F = 250mm, Thorlabs, and AC254-150-AB-ML F =150 mm, Thorlabs). The photon stimulation light combines with imaging light by a polarized beam splitter (CCM1-PBS252/M; Thorlabs) between tube lens and dichroic mirror.

The hologram is calculated by compressive sensing weighted Gerchberg–Saxton algorithm^41^ and loaded on SLM which controlled by a custom software on MATLAB. To compress the axial profile, a large beam diameter is applied to achieve an effective excitation NA (∼0.5). Then by generating a spot array on each target, the target neuron can be fulfilled with a square of 10-12 um diameter and with an average power density of 0.15-0.3 mW/µm^2^.

The spatial coordination between imaging light and photo stimulation light is calibrated before each optogenetics experiment. Two dark points are generated by photon bleaching a fluorescent microscope slide (FSK3, Thorlabs) for both imaging light and photo stimulation light. After calculating the displacement between them, a coordination shift is added to the hologram calculation.

During the two-photon optogenetics experiment, for each zebrafish larva, 6-8 sessions are conducted depending on the specific expression. For each session, a single pretectal *KalTA4u508* neuron is randomly selected according to mCherry expression and activated. Each session contains 5 trials, whereby each trial entails a 10-second duration of neuronal activation followed by a 30-second intertrial interval.

### Single-photon optogenetic stimulation

Zebrafish larvae were placed in the petri dish and fixed with agarose. The eye and tail are freed by carefully cutting the agarose. An optical fiber, 200 µm in diameter, is held 5mm above the zebrafish larvae and guided the light from a blue light fiber-coupled LED (M470F4, Thorlabs). The irradiance at zebrafish larvae is 12-20 mW/cm^2^ and adjusted by pulse-width modulation. Each trial contains a 2 s stimulation period and there is a 15 s interval between each trial. In total 60 trials are conducted for each fish. The whole petri dish is illuminated by an 850 nm LED array. Behavior is recorded at 100 Hz frame rate and analysed by custom Python software. The ChR2 expression is recorded after the experiment by two-photon imaging as a reference.

### Behavioral Tracking

We tracked behavior using ztrack, a Python-based custom software. For eye tracking, we applied image thresholding around the fish’s head, extracting the three largest contours from the image. The closest pair was identified as eyes, and the third as the swim bladder. Using the cv2.fitEllipse function from OpenCV, we fitted ellipses to these contours to find the centers and orientations of the eyes and swim bladder. We distinguished left from right eye based on the direction of the cross product between the vectors from the swim bladder to each eye. The fish’s midline was defined as the line from the swim bladder’s center to the midpoint between the eyes. The eye angles were computed as the angles between the major axes of the eyes and the midline. To track the tail, we used an iterative algorithm that traced 11 points along the tail, beginning at the most posterior point of the swim bladder. At each step, the algorithm identifies the next point on the tail by finding the brightest pixel among locations that were equidistant from the current point.

### Behavioral Data Analysis

Eye convergence events were detected as peaks in the convolution of gaussian filtered (σ = 0.04 s) vergence angle (L + R eye angle) with a step function (width 1 s), similar to^13^, and manually corrected. Tail bouts were detected by thresholding the rolling standard deviation of tail angle over a 50 ms window. The asymmetry of each bout was calculated by first integrating the curvature over the last quarter of the tail and then determining the fraction of time points with curvature integrals that matched the sign of the bout’s mean curvature.

### Processing of Calcium Imaging Data

For each imaging plane, the non-rigid motion correction was performed using suite2p^42^. In Figure 1E, pixel intensities were preprocessed by z-score normalization across frames prior to analysis. In Figure 1F, AF7 activity was calculated as the mean intensity of pixels within the manually annotated AF7 mask, followed by z-score normalization across time. The tuning curve in Figure 1G was derived from the peak normalized AF7 activity within the stimulus interval.

For neuron-based analyses, the locations of regions of interest (ROIs) and their ΔF/F were extracted using the CNMF algorithm from the CaImAn package^43^. The 10^th^ percentile of a rolling 3-minite time window was used as the baseline for calculating ΔF/F. ROIs exhibiting a signal-to-noise ratio (SNR), as computed by CaImAn, below 2.5 were excluded from further analysis. The ΔF/F for each ROI was normalized by subtracting the mean and then dividing by the standard deviation obtained across all imaging sessions, followed by smoothing with a Gaussian filter (σ = 0.33 s).

### Image Registration

Image registration was performed using the antsRegistration function from Advanced Normalization Tools (ANTs)^44^ using parameters described^45^. Anatomical z-stacks were registered to the templates provided by the mapZebrain atlas^29^.

### Assignment of Neurons to Anatomical Regions

For each region of interest (ROI), we calculated its x and y coordinates based on the center of mass of its spatial component. To determine the z coordinates, we identified the depth of each imaging plane by finding the z-stack layer that maximizes the Pearson correlation with the average fluorescence image of that plane. We then converted the 3D coordinates from the z-stack space to the atlas space using the function antsApplyTransformsToPoints. Lastly, we identified the anatomical region of each ROI by referencing annotated volume of the atlas at the transformed 3D coordinates.

### Spatial *p*-value Analysis

For all maps showing the anatomical locations of functionally classified neurons from multiple fish, we retain only the neurons that demonstrate spatial co-localization across fish, as determined through the spatial *p*-value analysis described^30^.

### Regression Analysis

We generated stimulus regressors by convolving boxcar functions indicating stimulus presence with a calcium impulse response function (CIRF), which was modeled as a difference-of-exponentials^46^. We conducted multiple linear regression analyses using sklearn.linear_model.LinearRegression. In the analysis in Figure 2, we used 12 stimulus regressors (−50°, …, −0°, +0°, …, +50°) to model the ΔF/F for each ROI. Separate regression models were used for different ocular conditions, yielding one R^2^ value per condition. For the analysis in Figure 3, we concatenated the ΔF/F for each neuron across all ocular conditions. This concatenated data was then modeled as a weighted sum of 38 regressors, composed of 12 stimulus regressors for each of the three ocular conditions (left-eye (LE), right-eye (RE), and binocular) plus 2 motor regressors. For the analysis in Figure S4C, we used 8 stimulus regressors corresponding to the 8 stimulus types. An intercept term was included in all regression models.

### Ocular Specificity Assignment

To categorize neurons into one of four ocular types in Figure 2, we first compute the R^2^ values for ipsilateral and contralateral conditions. For neurons in the left hemisphere, R^2^_ipsi_ refers to R^2^_LE_, and R^2^ refers to R^2^. For neurons on the right, these assignments are reversed. Neurons are then classified as follows: binocular neurons: both R^2^_ipsi_ and R^2^_contra_ ≥ 0.3; mono-ipsi neurons: R^2^_ipsi_ ≥ 0.3 and R^2^_contra_ < 0.3; mono-contra neurons: R^2^_contra_ ≥ 0.3 and R^2^_ipsi_ le < 0.3. Neurons that do not fit any of these criteria are classified as “Other”.

### Hierarchical Clustering

In Figure S4, we clustered neurons by their responses to the eight stimulus types using affinity propagation followed by hierarchical clustering, as descripted^26^.

### Analysis of Imaging Data from Optogenetics Experiments

For the analysis in Figure 5D, neurons were categorized as *lhx9:Gal4* or *KalTA4u508* depending on which corresponding z-scored fluorescence z-stack exhibited a higher z-scored fluorescence intensity at the neuron’s location.

### Quantification and Statistical Analysis

For each tuning curve, we determined the 95% confidence intervals through bootstrapping. We applied the Bonferroni method to adjust the *p*-values for multiple comparisons. For the box plots, the box spans from the first to the third quartile of the data, with the median marked by a line inside the box. The caps of the whiskers mark the most distant data points that are no more than 1.5 times the interquartile range away from the box. To estimate the 95% confidence intervals, we generate 1,000 bootstrap samples by resampling the data points with replacement. We then calculated their means and computed the 2.5^th^ and 97.5^th^ percentiles of these means.

## Data and materials availability

All data are available in the main text or the supplementary materials. Code is available at https://github.com/SemmelhackLab/binocular-paper

## Acknowledgments

Funding was provided by the Hong Kong Research Grant Council to J.L.S. (16103522, 16101221, 16103118, and 26100617) and J.Y.Q. (16102122) and the Chau Hoi Shuen Foundation. We thank Aristides Arrenberg and members of the Semmelhack lab for feedback on the manuscript.

## Author contributions

Conceptualization, G.T. Methodology, G.Y. and Y.H. Software, G.Y. and T.K.C.L. Formal Analysis, T.K.C.L Investigation, G.T., T.K.C.L. and G.Y. Resources, G.Y. and B.K. Writing – Original Draft, G.T. and J.L.S. Writing – Review & Editing, all authors. Supervision, J.L.S. and J.Y.Q. Funding Acquisition, J.L.S. and J.Y.Q.

## Supplemental information

Document S1. Figures S1–S8.

Movies S1 to S2.

## Supplemental Video Legends

**Supplemental Video 1: Example trials of 2-photon calcium and behavioral recordings.** Left: ΔF/F signal of cell bodies in the imaging volume, from coronal, dorsal, and sagittal views. Right: prey stimulus and behavioral recording. Imaging speed was 3 volumes per second, and behavior was recorded at 200 frames per second. Playback has been speed up 4x. 1. Both eyes exposed condition 2. Right eye exposed condition 3. Left eye exposed condition. Larvae expressing nuclear localized GCaMP6s (*Tg(elavl3:Hsa.H2B-GCaMP6s)jf5Tg*) were used for these imaging experiments.

**Supplemental Video 2: Examples of prey capture bouts evoked by optic fiber.** Top: eye and tail angle plots over time. Bottom: behavioral video with eyes and tail tracked. Larvae were *Tg(KalTA4u508;lhx9Gal4;UAS-GCaMP6s;UAS:ChR2(H134R)-mCherry)*.

