## Supplemental Figures for "Binocular integration of prey stimuli in the zebrafish visual system"

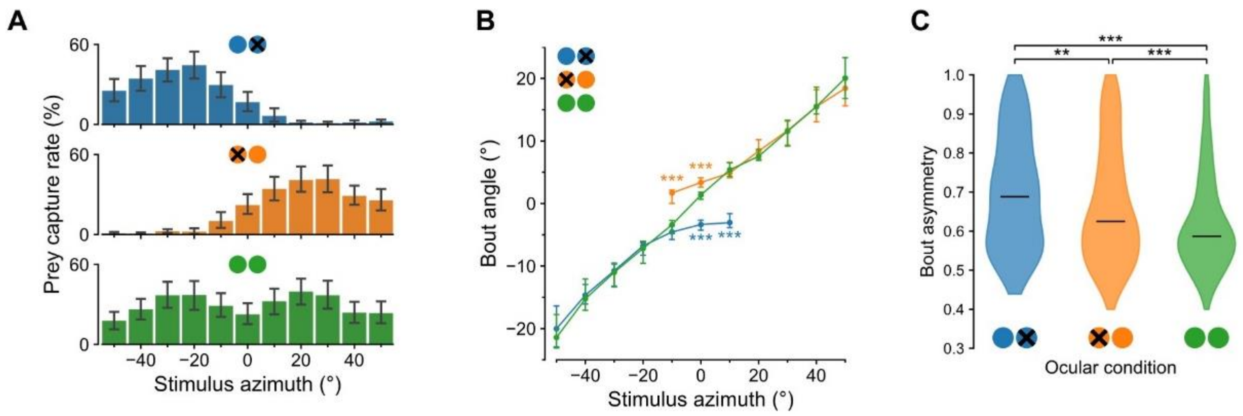


**Figure S1: Occlusion of one eye affects prey capture behavior, related to Figure 1.** (A) Prey capture response rates (percentage of trials with eye convergence) over 11 azimuths in each of the ocular conditions. Error bars indicate 95% confidence intervals. n = 41 fish. (B) Bout angle (maximum angle of the first half beat) during prey capture in response to the stimuli in different ocular conditions. Dots and error bars indicate the median and 95% confidence interval, respectively (Kruskal-Wallis test followed by Dunn's test, *** P < 0.001, otherwise P > 0.05). (C) Bout asymmetry (fraction of time curvature direction matched direction of initial turn) during hunting bouts (defined by eye convergence) in response to prey at 0° in different ocular conditions. Horizontal lines indicate the median. n = 282, 396 and 437 bouts from 41 fish for left-eyed, right-eyed and binocular conditions, respectively (Kruskal-Wallis test followed by Dunn's test, ** P < 0.01, *** P < 0.001).

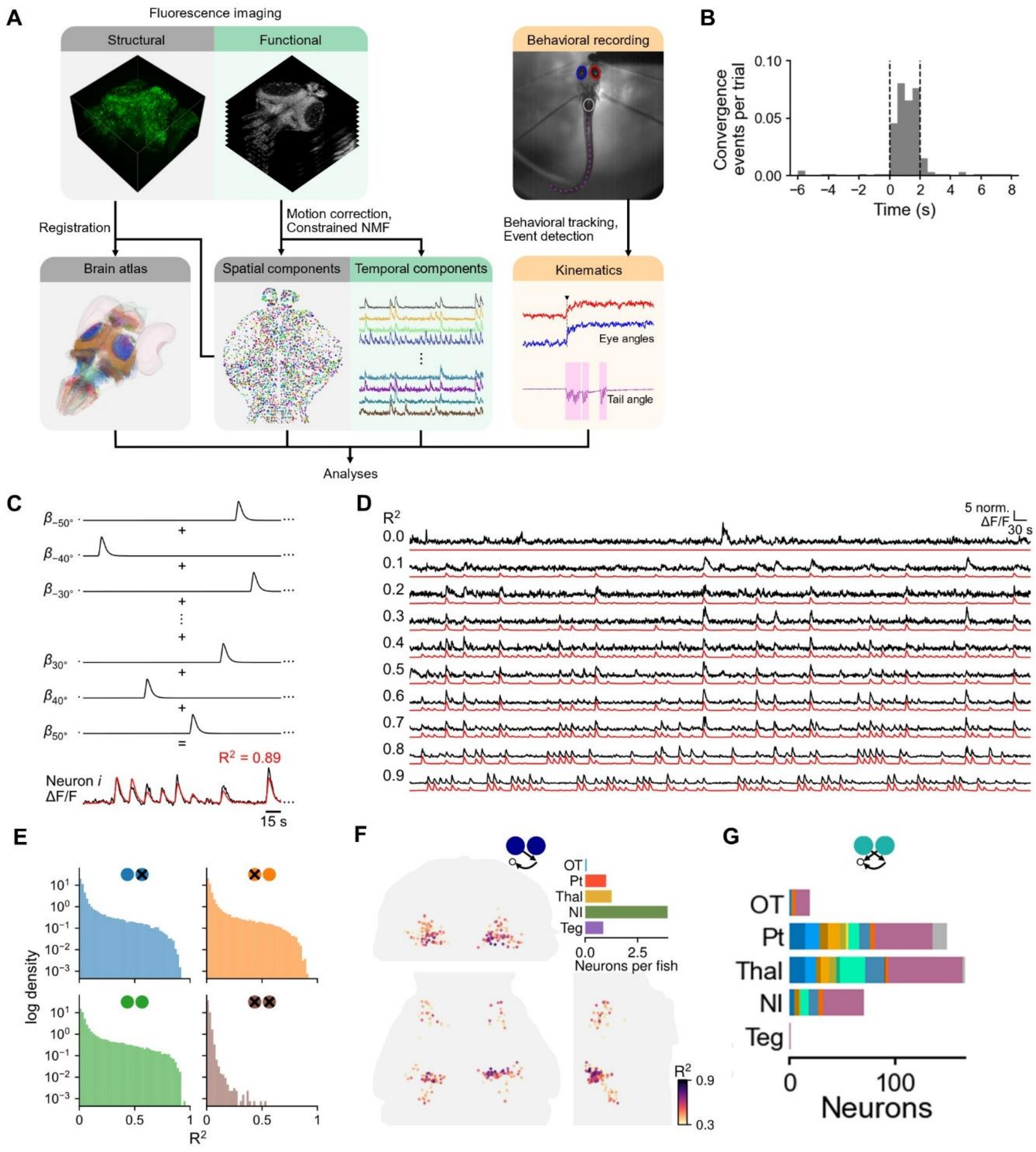


**Figure S2: Analysis pipeline and R2 threshold for PRNs, related to Figure 2.** (A) Schematic of integrated pan-neuronal 2-photon calcium imaging and behavioral recording pipeline. (B) Timing of convergence events within 0° prey trials in the binocular condition. Dashed lines represent stimulus interval. (n = 15 fish). (C) Schematic of linear regression model. Regressors corresponded to the different prey stimuli (azimuths -50° left to 50° right). Red line is predicted activity based on the regression model of that neuron. (D) Calcium traces of example neurons with R2 ranging from 0 to 0.9. Red line is predicted activity based on the regression model of that neuron. (E) R2 distributions of neurons in each ocular condition (n = 15 fish). (F) Anatomical locations of monocular ipsi PRNs (mono-ipsi). (G) Number of bino-PRNs in each brain area over the 15 fish in the dataset. Each color represents one fish.

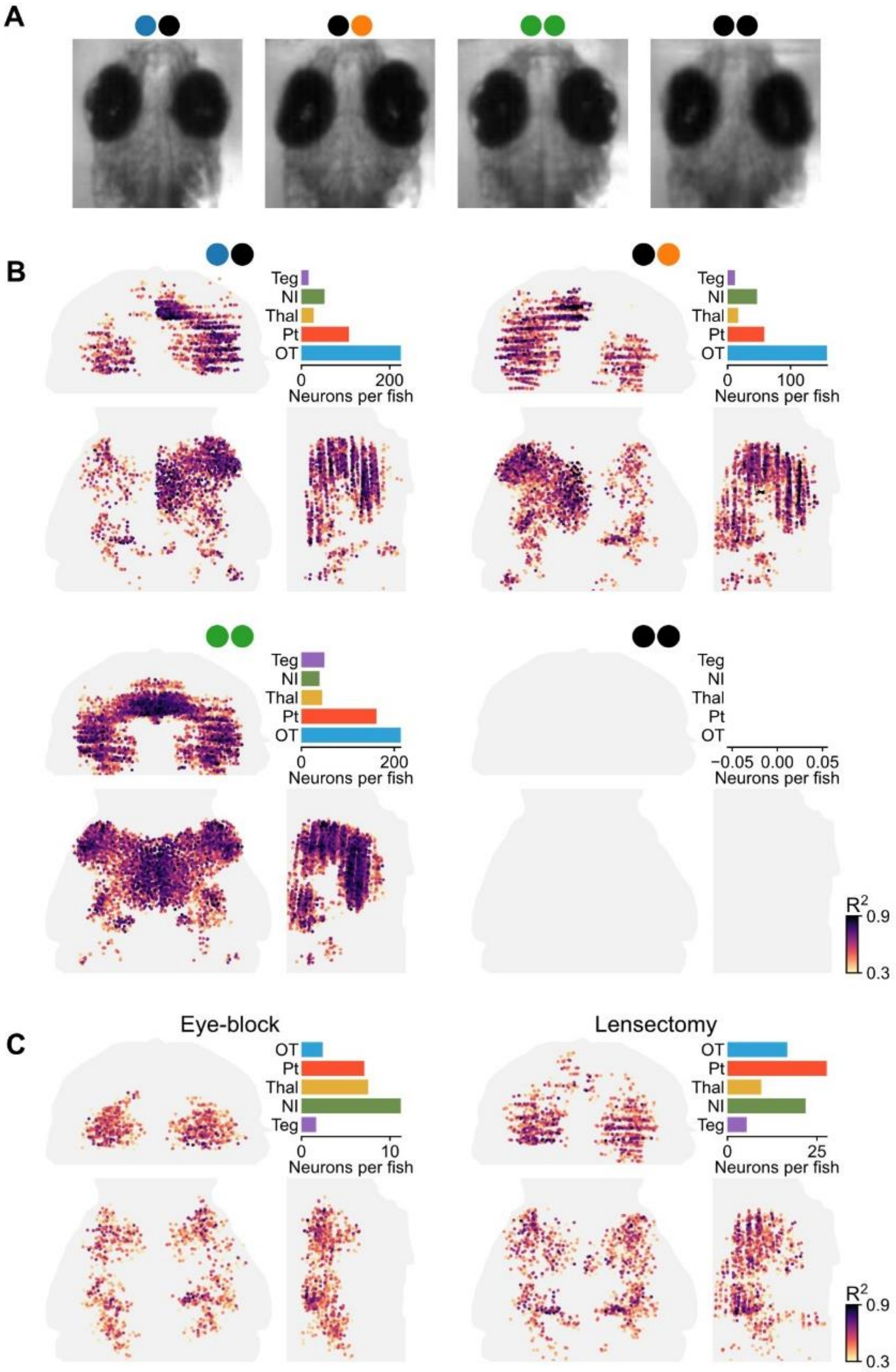


**Figure S3: Lesectomy of one eye results in the same pattern of PRN activation as blocking one eye, related to Figure 2.** (A) Monocular left, monocular right, sham, and bilaterally lensectomized larvae. (B) Maps of PRNs in different lens removal conditions (n= 4 for left-eyed, 7 for right-eyed, 7 for sham, 2 for bilateral lensectomy). (C) Comparison of ipsi PRN (PRNs activated by the ipsilateral eye) position in monocular eye-blocked and monocular lensectomized larvae (left and right-eyed combined).

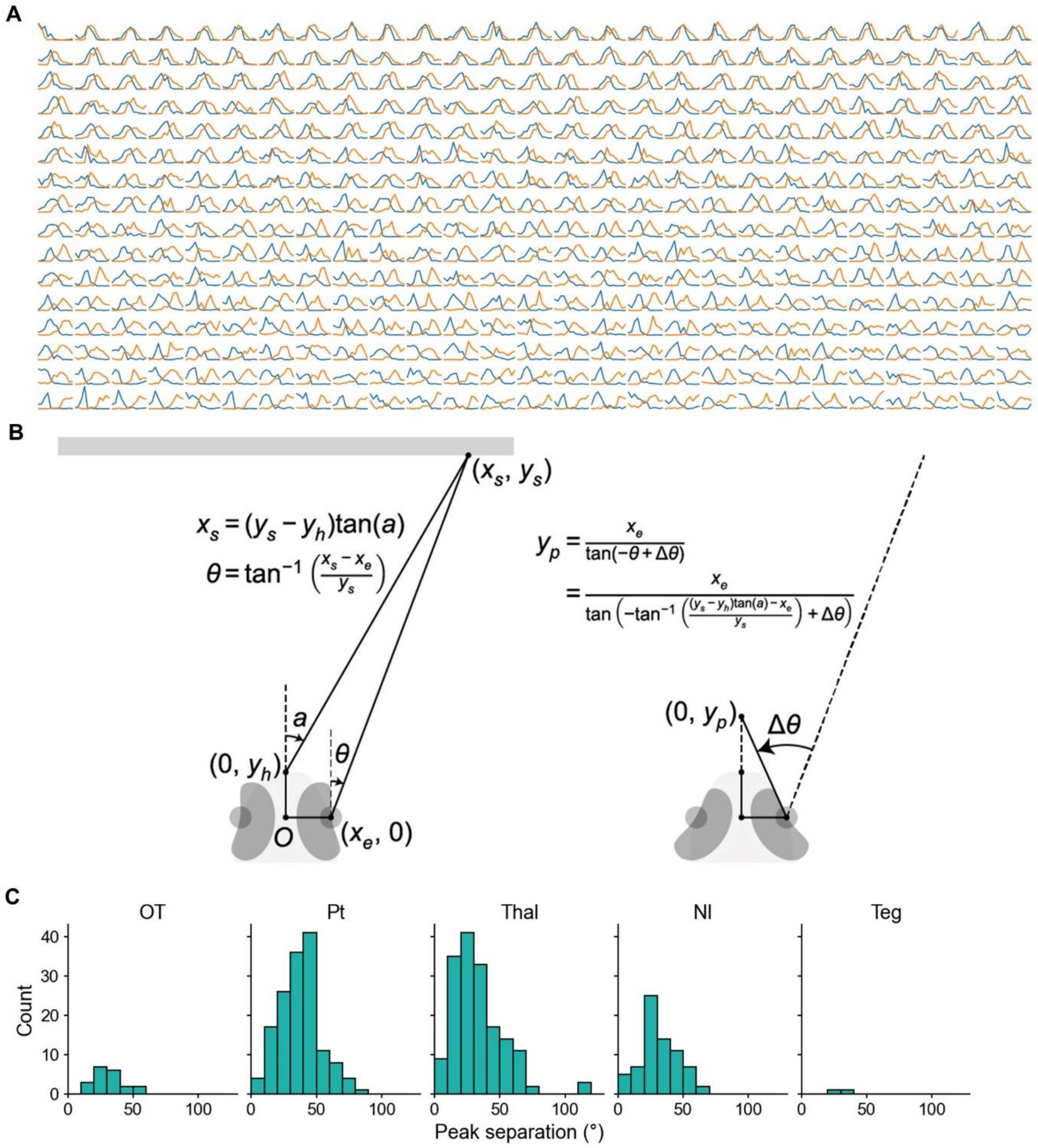


**Figure S4: Bino-PRN tuning curves, calculation of prey distance from peak separation, and distribution of peak separations in different brain areas, related to Figure 4.** (A) Left eye (orange) and right eye (blue) tuning curves for all bino-PRNs, for prey stimuli presented at azimuths from -50° to +50°. (B) To calculate the distance of a prey object ($y_p$) that would activate a bino-PRN with a particular left eye-right eye peak separation, we first calculate the angles *a* and θ (the angle of the visual line with respect to the vertical axis) for a given stimulus location $x_s$, and a screen distance of 1 cm. We then calculate the distance of the prey object that would activate the same neuron after eye convergence by taking the change in eye angle as Δθ. (C) Distributions of left eye-right eye peak separations for bino-PRNs in the five brain areas containing bino-PRNs.

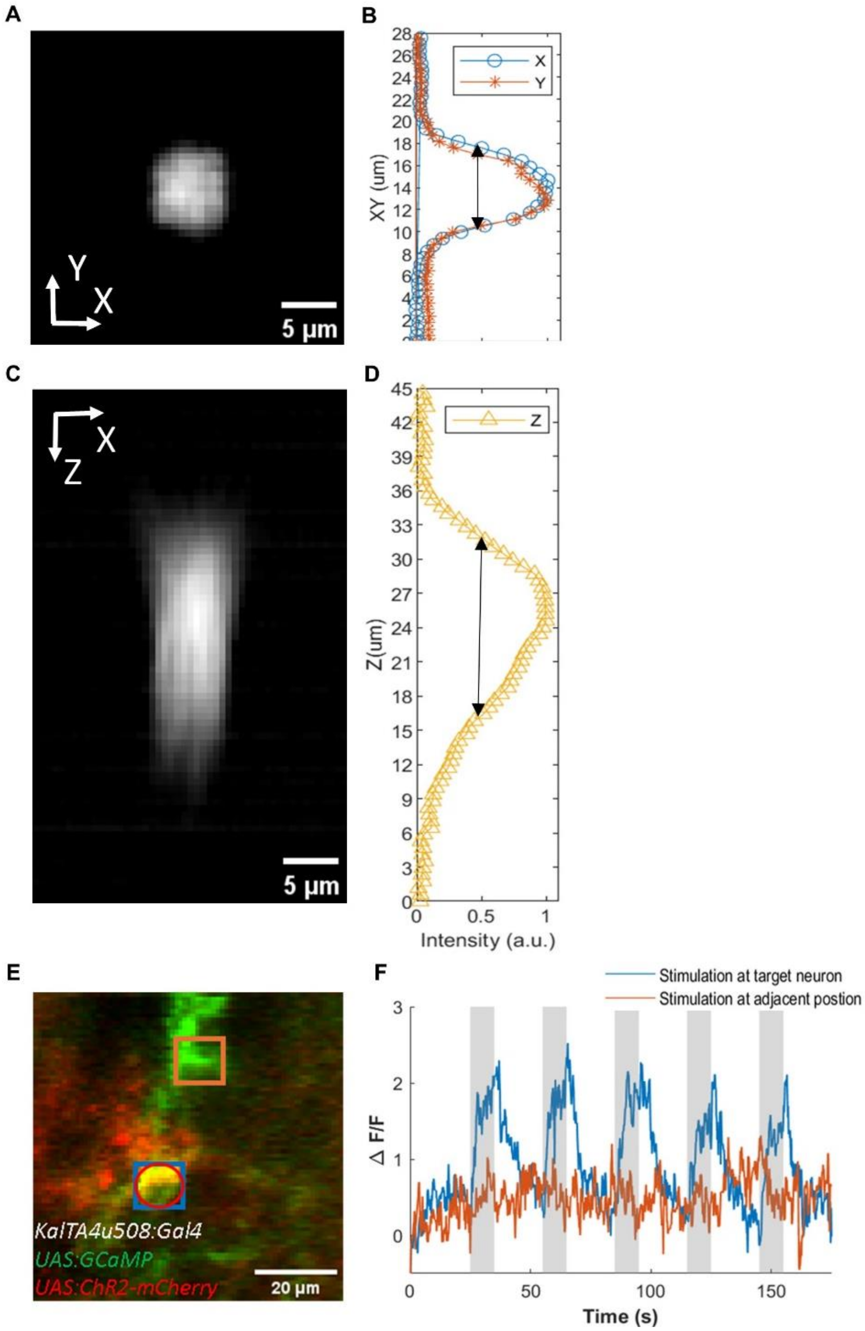


**Figure S5: Resolution and efficacy of holographic optogenetic activation, related to Figure 7.** (A) 3D holographic patterns, projected in X and Y, achieved by reversing the image bleached in a fluorescein microscope slide. (B). Profile of the bleached area (normalized), (Full width at half maximum (FWHM): 7.2 µm (X) and 6.5µm (Y)). C. 3D holographic pattern projected in the Z direction and (D). its profile (normalized), (FWHM: 16.7 µm (Z)). (E) The regions selected for testing targeting accuracy in Tg(KalTA4u508:Gal4;UAS:GCaMP6s; UAS:ChR2-mCherry) larvae. Blue square: excitation position (at the target neuron) on a u508 APN neuron. Orange square: adjacent ROI. Green: GCaMP; Red, ChR2-mcherry. (F) The calcium trace of the target neuron (blue) and adjacent neuron (orange) during two-photon optogenetic excitation periods (grey shading).

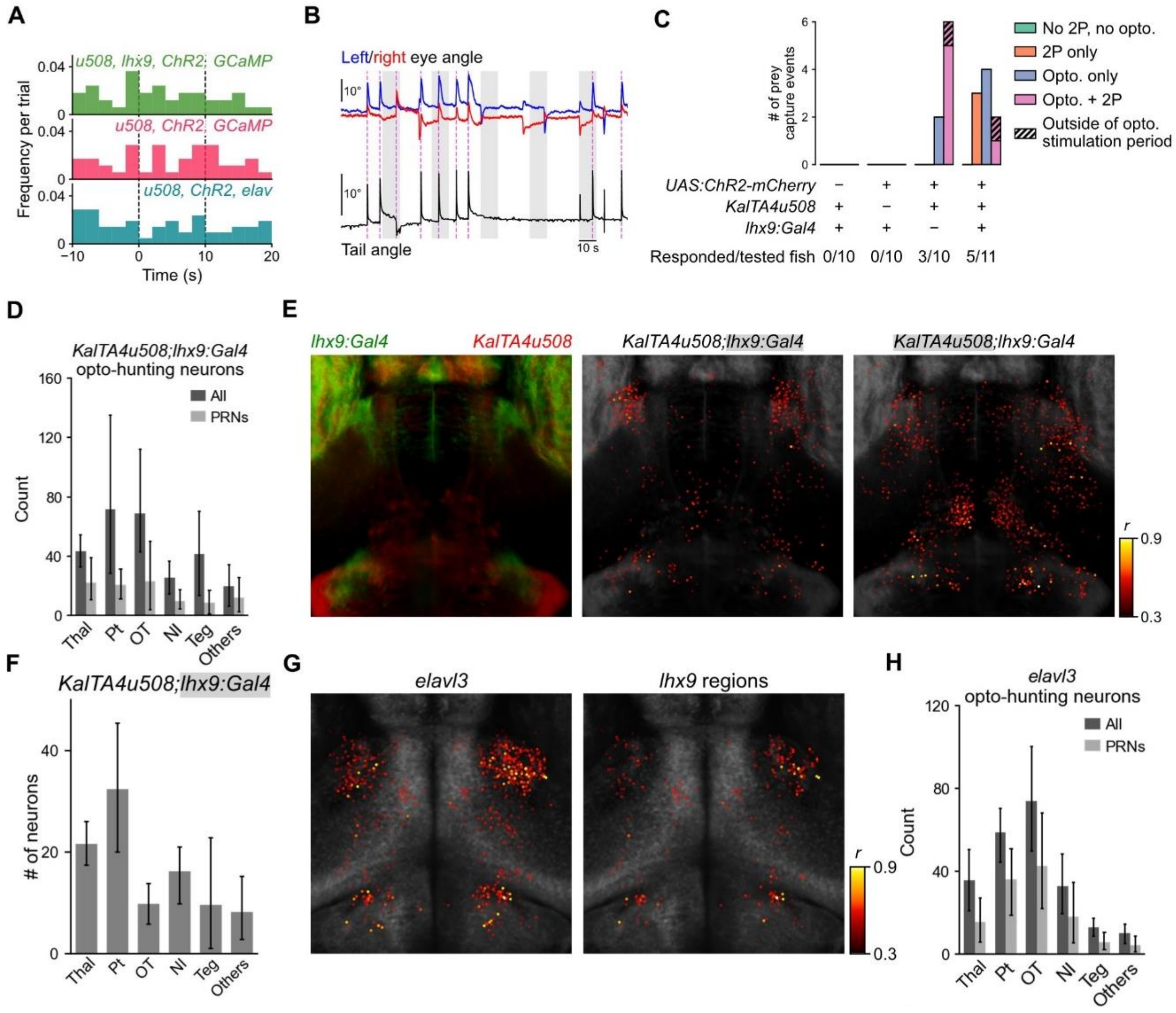


**Figure S6: Opto-hunting behaviour requires ChR2 expression in command-like neurons and is correlated with *lhx9+* neurons and neurons within the DT, PT and NI in a panneuronal line, related to Figure 7.** (A) Distributions of hunting onset time during opto/imaging sessions. (B) Eye convergence and tail movements triggered by optogenetic activation of *KalTA4u508* APN neurons. (C) Total number of hunting initiations over all fish of each genotype triggered by scanning and activation. Each condition was tested ten times per fish. (D) Average number of opto-hunting neurons in the *lhx9 and KalTA4u508* fish (dark grey) and number of PRNs within that population. Error bars indicate ±1 SD. n = 5 fish. (E) Left: the *lhx9* (green) and *u508* (red) masks. Middle: opto-hunting neurons within the *lhx9* mask. Right: opto-hunting neurons in the *u508* mask. Dorsal view. n = 5 fish. (F) Average number of opto-hunting within the *lhx9* mask. (G) Distribution of opto-hunting neurons labelled by *elavl3* after optogenetic activation of *KalTA4u508* APN neurons. Left: all *elavl3* opto-hunting neurons. Right: *elavl3* opto-hunting neurons within the *lhx9* mask. (H) Average number of total *elavl3* opto-hunting neurons in each fish, and number of PRNs within that population. Error bars indicate ±1 SD. n = 5 fish.

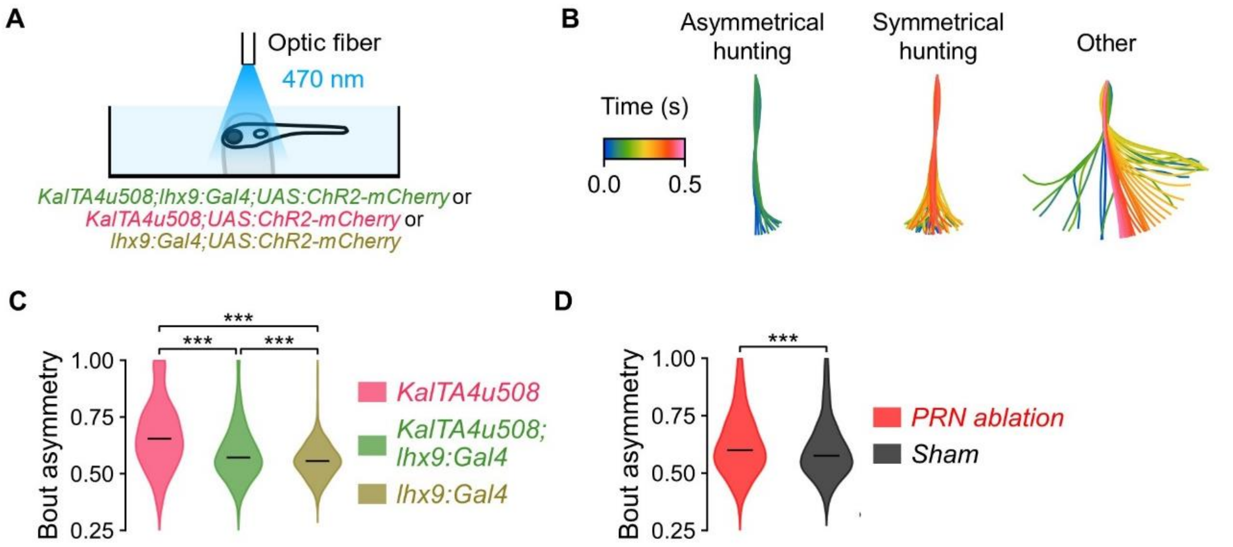


**Figure S7: Activation of lhx9+ neurons evokes opto-hunting bouts, related to Figure 7.** (A) Schematic of the optogenetic activation experiment with larvae bearing one or both gal4 lines. (B) Examples of the asymmetric and symmetric hunting bouts (defined by eye convergence) and other non-hunting bouts evoked by optic fiber stimulation. (C) Bout asymmetry of opto-hunting bouts (defined by eye convergence) in the three genotypes (Kruskal-Wallis test followed by Dunn's test, *** $P < 0.001$). (D) Asymmetry of opto-hunting bouts of KalTA4u508; lhx9:Gal4; UAD:ChR2-Cherry larvae after ablation of ~20 PRNs in the NI, or the same number of non-PRNs at a similar depth for the sham condition.
